# *Shigella*’s c-di-GMP specific PDEs Modulate Biofilm and Virulence Phenotypes

**DOI:** 10.64898/2026.06.22.733758

**Authors:** Candice N. Churaman, Benecia Angelica, Andrew W. Thompson, Benjamin J. Koestler

## Abstract

To establish infection and cause disease, the intracellular pathogen *Shigella* must successfully navigate a series of host defenses and distinct microenvironments within the human body. One way *Shigella* navigates these enviroments is by using the secondary messenger c-di-GMP, which regulates many different bacterial behaviours. C-di-GMP is synthesized by diguanylate cyclases (DGCs) and broken down by c-di-GMP specific phosphodiesterases (PDEs). In this study, we investigated how *Shigella*’s c-di-GMP specific PDEs impact c-di-GMP turn-over and subsequently biofilm and virulence phenotypes. We knocked out each of *Shigella’s* six c-di-GMP specific PDEs to determine how these PDEs impact biofilm, virulence and c-di-GMP levels within the bacterial cell. We found that these PDEs negatively regulate c-di-GMP levels while modulating *Shigella*’s virulence and biofilm behaviour. We also noted that altering expression of these *Shigella* PDEs changes bacterial cell size. Transcriptome analysis revealed that a *Shigella* Δ*pdeB* strain showed reduced expression of many genes, including the virulence genes *ipgD* and *ipgE*, as well as genes associated with lipid metabolism. We confirmed that a *Shigella* Δ*pdeB* strain had altered levels of stearic acid, and expression of *pdeB* alters *Shigella* antibiotic susceptibility. This study highlights the complexities of c-di-GMP signaling in regulating numerous *Shigella* pathways.

## Introduction

*Shigella*, an enteric pathogen belonging to the *Enterobacteriaceae* family, is the sole etiological agent of shigellosis, which causes bloody diarrhea, fever and abdominal cramps (1–3). *Shigellosis* outbreaks often occur in developing nations with less advanced sanitation infrastructure, disproportionately impacting children under the age of 5 (3–5). *Shigella* infects approximately 125 million persons globally every year (6). While Shigellosis mortality rates have modestly declined over recent years, the emergence of multi-drug resistant strains has been increasing exponentially (7). There are no available licensed vaccines for protection against *Shigella* infection (8,9). As such, there is a need to identify novel drug targets to combat this disease (10,11).

*Shigella* directly descended from commensal *Escherichia coli,* and *Shigella* species still share a high level of genetic similarity; for example, *S. flexneri* 16S rRNA sequence is >99% identical to *E.coli*, and the overall genetic divergence is approximately 1.5% (12–14). Though being so closely related, *Shigella* has both lost and gained functions in the process of pathoadaptation, making it distinct from *E.coli* (15). For example, *Shigella* has gained a virulence plasmid and pathogenicity islands, and lost motility and the ability to decarboxylate lysine (16,17). It is these loss and gain of functions which has allowed *Shigella* to thrive successfully as an intracellularly pathogen (18).

*Shigella* withstands the various physical and chemical barriers posed by the human gastrointestinal tract in order to survive and replicate, including chemical barriers of the digestive track such as the acidic environment of the stomach, and bile in the small intestines (7,19–22). *Shigella* is also able to overcome the physical barriers posed by the colonic epithelium to invade and replicate (12). Here, *Shigella* is engulfed by specialized epithelial cells known as microfold cells (M-cells) in Peyers patches (12). M-cells transcytose *Shigella* to the intra-epithelial pocket which contains macrophages and lymphocytes, that phagocytose the bacterial cells (12,23); *Shigella* is able to evade the lethal effects of these macrophages by inducing pyroptotic cell death of the macrophage (24). After, *Shigella* invades colonic epithelial cells from the basolateral surface using a type 3 secretion system (T3SS) (25). Once inside a host cell, *Shigella* replicates to high cell density, and then exploits host cytoskeleton synthesis using the protein IcsA to initiate actin-based motility, propelling the bacterium into a neighboring cell (21,26,27).

To survive in the human body, bacteria like *Shigella* alter their behavior by detecting and responding to specific environmental stimuli. One way bacteria relay these signals is by using second messengers like cyclic dimeric guanosine-monophosphate (c-di-GMP)(28,29). Enzymes that synthesize and break down c-di-GMP typically have N-terminal sensory domains that interact with environmental ligands, which then alters the activity of C-terminal domains that synthesize or hydrolyze c-di-GMP, which in turn regulates gene expression or protein function (29–33). Diguanylate cyclases (DGCs) synthesize c-di-GMP from GTP, and c-di-GMP specific phosphodiesterases (PDEs) hydrolyze c-di-GMP (32,34). PDEs contain either an EAL domain that can break down c-di-GMP into 5’-phosphoguanylyl-(3’-5’)-guanosine (pGpG) or they can contain a HD-GYP domain which can hydrolyze c-di-GMP into 2 molecules of GMP (30,31,34).

Compared to *E.coli*, *Shigella* has lost majority of its c-di-GMP turn over enzymes. In our previous studies, we characterized *S. flexneri*’s DGCs and found that four *S. flexneri* DGCs (DgcF, DgcC, DgcI and DgcP) produced c-di-GMP, and disruption of these genes altered acid resistance, biofilm formation, and virulence phenotypes (35,36). Specifically, all four DGCs modulated *S. flexneri* biofilm formation (36), Δ*dgcF* and Δ*dgcC* mutant strains demonstrated invasion defects, and a Δ*dgcF* mutant was deficient in its ability to form plaques in Henle-407 cells, and exhibited increased survival when acid shocked (36). From these findings, we concluded that c-di-GMP signalling is playing a role in regulating *Shigella*’s behaviour associated with human colonization. *Shigella’s* genome has 6 intact c-di-GMP PDEs and another 6 c-di-GMP PDEs annotated as pseudogenes due to the presence of either a non-sense or frameshift mutation. *Shigella* have lost an additional other 5 c-di-GMP PDEs. To date there have been no studies on *Shigella*’s PDEs.

Because *Shigella*’s DGCs regulated c-di-GMP levels and various phenotypes, we decided to investigate *Shigella*’s 6 intact c-di-GMP PDEs (*pdeB, pdeD, pdeG, pdeH, pdeK* and *pdeN*) to determine how these non-pseudogene PDEs are able to modulate *Shigella*’s biofilm and virulence phenotypes and c-di-GMP levels. We found that *pdeB, pdeG, pdeH, pdeK* and *pd*e*N* contributes to *Shigella*’s biofilm formation. We also found that the *Shigella* Δ*pdeB* and Δ*pdeG* mutant strains showed increased invasion of Henle-407 cells. *Shigella*’s Δ*pdeG* and Δ*pdeN* had increased plaque sizes while Δ*pdeK* had a decreased plaque size. When we measured c-di-GMP, we found that the *Shigella* mutant Δ*pdeB*, Δ*pdeD* and Δ*pdeH* strains showed increased c-di-GMP levels. We then investigated how *pdeB* influences *Shigella’s* gene expression. We observed that mutation of *pdeB* alters expression of genes associated with various pathways including virulence and lipid metabolism, which we confirmed by assessing changes in the lipid profile. Together, this work demonstrates that *Shigella*’s PDEs contribute to regulating c-di-GMP and subsequent behavioural changes through altering gene expression.

## Results

### *S. flexneri* encodes 6 PDEs homologous to *E. coli*

*E.coli* MG1655 encodes 29 c-di-GMP turnover enzymes in its genome, 12 of which are DGCs, 10 are PDEs and 7 are hybrid proteins that contain both a GGDEF and an EAL domain. In contrast, *Shigella flexneri* 2457T encodes 10 c-di-GMP turnover enzymes that are at least 97.9% identical at the nucleotide level to those encoded by *E.coli* MG1655. These include 4 DGCs, 5 PDEs containing an EAL domain, and 1 hybrid protein with a degenerate GGDEF domain and an active EAL domain. Of note, neither *E. coli* nor *S. flexneri* encode any HD-GYP domain containing genes. The remaining 19 *S. flexneri* genes encoding c-di-GMP turn-over proteins are annotated as pseudogenes or have been deleted from the genome.

The *S. flexneri* intact PDE genes (genes that do not carry nonsense or frame shift mutations) homologous to *E. coli* are *pdeB*, *pdeD*, *pdeG*, *pdeH*, *pdeK* and *pdeN* (Fig 1, Supplemental Table 1). *pdeB*, *pdeD*, *pdeG* and *pdeN* all share a similar structure; these genes encode a putative N-terminal CSS motif sensory domain, flanked by hydrophobic residues that are predicted to be transmembrane segments (29,37,38). In the C-terminal portion of these enzymes is a putative cytoplasmic EAL domain that grants c-di-GMP hydrolysis activity (37–40). *pdeK* encodes an N-terminal HAMP domain which relays an unknown signal to a putative periplasmic GAPES3 sensory domain); *pdeK* also encodes a putative cytoplasmic GGDEF-EAL domain. Of note, PdeK is predicted to be catalytically inactive in its c-di-GMP synthesis, as the active site motif encodes SGYDF in lieu of GGDEF (38,43). In *E. coli*, PdeK negatively regulates cellulose synthase (38,42). *pdeH* does not contain any predicted sensory domains or transmembrane domains (44); in *E. coli*, PdeH has been described as a PDE that maintains baseline c-di-GMP levels low in the cell (38,44).

**Figure 1.**
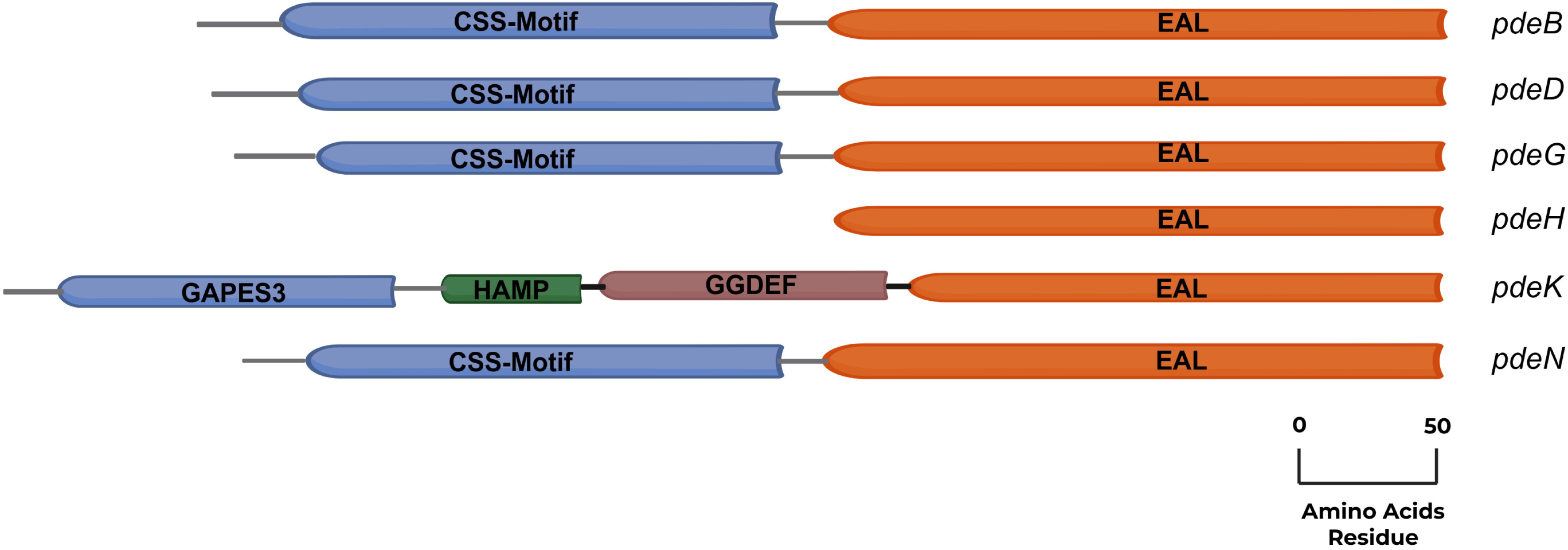
Schematic diagram of *Shigella*’s 6 intact PDEs was determined using UniProt (45). N-terminal to the cytoplasmic EAL domains (orange) are the periplasmic sensory domains (purple). On either side of the periplasmic sensory domains are predicted transmembrane segments (grey).

### *Shigella*’s PDEs do not impact growth

To determine if *S. flexneri* PDEs contribute to regulating behaviors associated with human colonization, we created *S. flexneri* strains where *pdeB*, *pdeD*, *pdeG*, *pdeH*, *pdeK* and *pdeN* were each deleted by homologous recombination (46). We also cloned each of these genes on a plasmid, under the control of an IPTG inducible promoter. If *S. flexneri* PDEs hydrolyze c-di-GMP, we hypothesize that knocking out *S. flexneri’s* PDEs will increase c-di-GMP levels, whereas expressing them from a plasmid will reduce c-di-GMP levels. C-di-GMP is a global regulator that influences many bacterial functions and is intertwined with many regulatory pathways (47). In *E.coli*, c-di-GMP regulates growth; *DgcN* plays a dual role in increasing levels of c-di-GMP and interfaces with the Z-ring, which results in cell division arrest (48). Notably, *dgcN* is a pseudogene in *S. flexneri* (*35,49*). Because c-di-GMP alters growth in *E. coli* and other bacteria, we first wanted to determine if *Shigella*’s PDE’s impact its growth. We performed growth curve analysis on both PDE mutant and plasmid expression strains. We monitored growth of each strain over a 24-hour period. In a minimal, defined growth medium (M63), our *S. flexneri* PDE mutant strains did not show significant differences in growth dynamics or growth rate, compared to WT (Fig. 2A & B). Likewise, when we expressed PDEs in their respective mutant strains, we did not observe any significant effects on growth dynamics or growth rate during the exponential phase when compared to WT carrying an empty plasmid (Fig S1A & B). We conclude that *S.flexneri* PDEs do not impact growth under the growth conditions used here.

**Figure 2.**
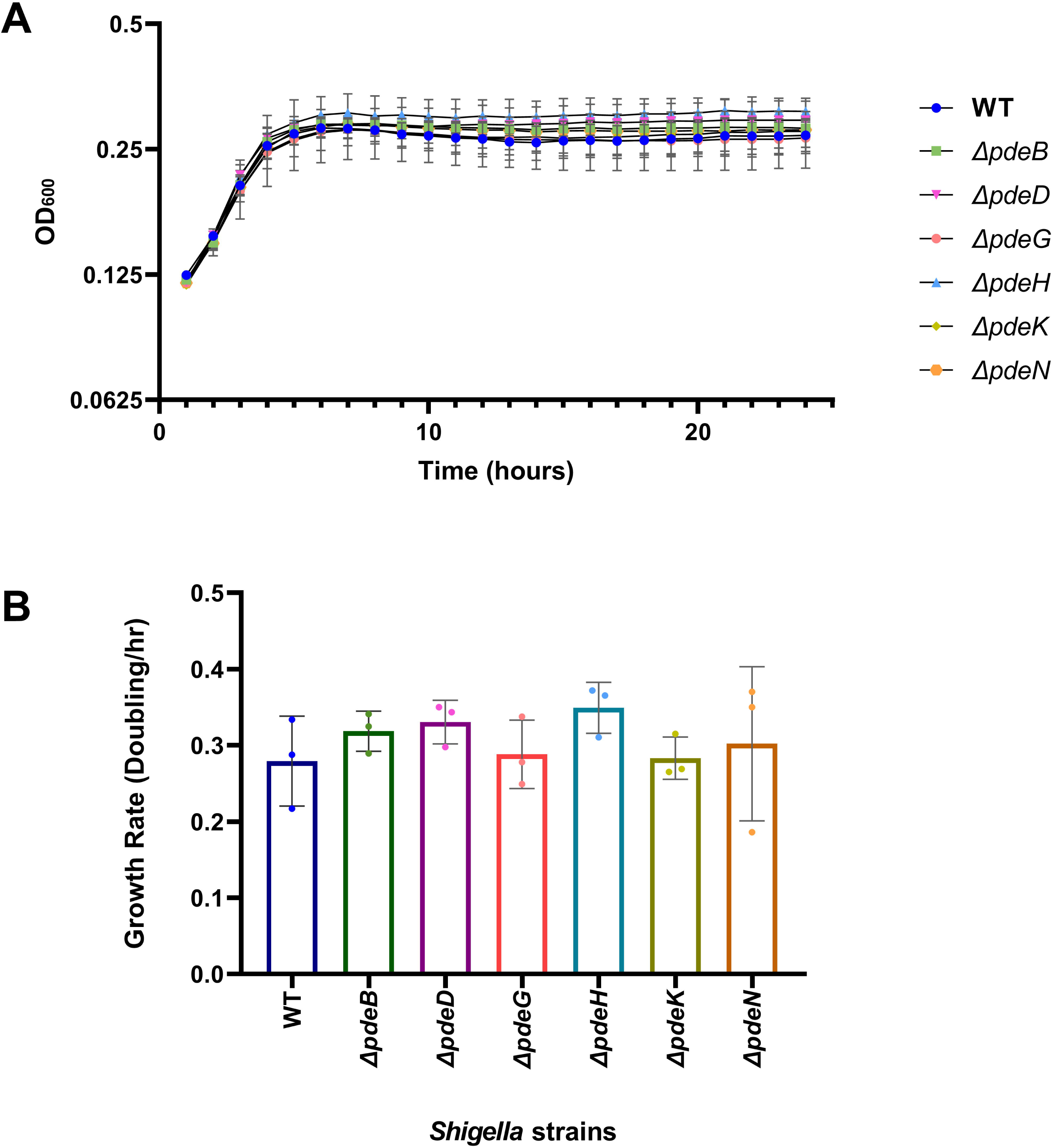
*Shigella’*s PDEs do not affect growth. (A) Growth curves were measured for *Shigella* mutant PDE strains over a 24 hours period to determine if the absence of any of these genes affected its growth. All strains grew similar to WT during the lag, log and stationary phase with no statistical difference, determined by Two-Way ANOVA with Dunnett’s multiple comparison test (p < 0.05) to compare WT to all mutant strains. (B) Growth rate was calculated during log phase growth to ensure each strain was growing at the same rate (B). There was no statistical difference between WT and the mutant strains’ growth rate. Statistical analysis was performed using a One-way ANOVA with Dunnett’s multiple comparison test (p <0.05) to compare WT to each mutant strain. Both panel A and B are a combination of three independent trials. Error bars indicate standard of deviation (SD).

### *S. flexneri* PDEs regulate biofilm formation

Knocking out *S. flexneri* DGCs reduces biofilm formation (36). To determine if *S. flexneri* PDEs regulate biofilm formation, we measured the biofilm formation of our *S. flexneri* PDE mutant strains using a biofilm micro-titre assay. We supplemented the growth medium with deoxycholic acid (DOC) and glucose, to induce biofilm formation (50). We included a *S. flexneri* Δ*bcsE* strain as a negative control, as *BcsE* is a positive regulator of cellulose synthesis, and both *E. coli* and *S. flexneri* Δ*bcsE* strains are defective in biofilm formation (36,51). We observed that *S. flexneri* mutant Δ*pdeB,* Δ*pdeG,* Δ*pdeH,* Δ*pdeK* and Δ*pdeN* strains showed a modest increase in biofilm formation compared to WT, while the mutated Δ*bcsE* strain had reduced biofilm formation compared to WT (Fig 3). As we expected, there was a statistically significant reduction in biofilm formation when each *S. flexneri* PDE was expressed in its respective mutant strain (Fig S3). Together, these data indicate that *S. flexneri* PDEs are regulating biofilm formation.

**Figure 3.**
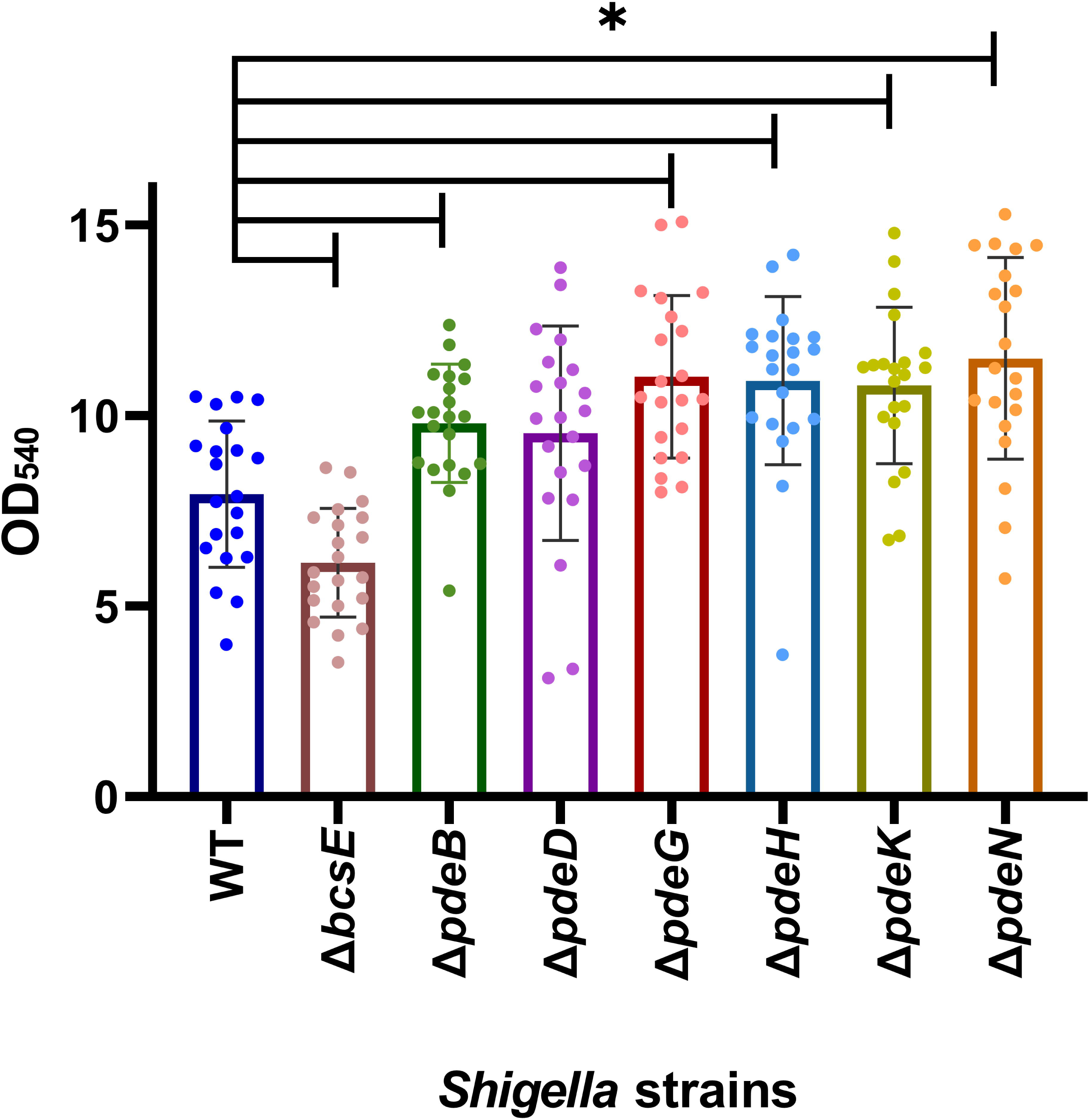
*S.flexneri* mutant PDE strains result in an increase in biofilm formation. A biofilm microtiter assay was performed using WT and *S.flexneri* mutant PDE strains. Δ*bcsE* was used as a negative control due to it having a biofilm defect (36). Bars indicate the mean OD_540_ while error bars indicate SD. Statistical analysis was perform using One-Way ANOVA with Dunnett’s multiple comparison tests (p < 0.05), comparing each strain to WT. * denotes statistical significance to WT.

### *S. flexneri pdeB* and *pdeG* regulate invasion

As part of its pathogenesis, *S. flexneri* invades colonic epithelial cells (12,24). We previously showed that altering *S. flexneri* c-di-GMP levels by expressing or deleting DGCs alters host invasion (36). It has also been demonstrated that deletion of *yfiB* in a different *S.flexneri* strain (Y394) reduces c-di-GMP levels, corresponding with reductions in adherence and invasion (52). To establish if *Shigella*’s PDEs are able to regulate host cell invasion as well, we used a Henle-407 tissue culture model of infection (53). We measured host cell invasion frequency of each of our 6 *S. flexneri* PDE mutant strains. WT was included as positive control, and we also included the strain CFS100 as a negative control; CFS100 is a *S. flexneri* strain lacking the virulence plasmid and thus incapable of invasion (54,55). We observed that the *S. flexneri* mutant Δ*pdeB* and Δ*pdeG* strains showed increased invasion frequency, with *S.flexneri* Δ*pdeB* have more than a 2-fold increase in host cell invasion compared to the WT strain (Figure 4A). *S.flexneri* mutated Δ*pdeD*, Δ*pdeH* Δ*pde*K and Δ*pdeN* strains did not have any effect on host cell invasion. The respective complementation strains for all mutant strains showed reduced invasion (Fig S2A). These data together indicate that *S.flexneri pdeB* and *pdeG* contribute to regulating invasion of Henle-407 cells.

**Figure 4.**
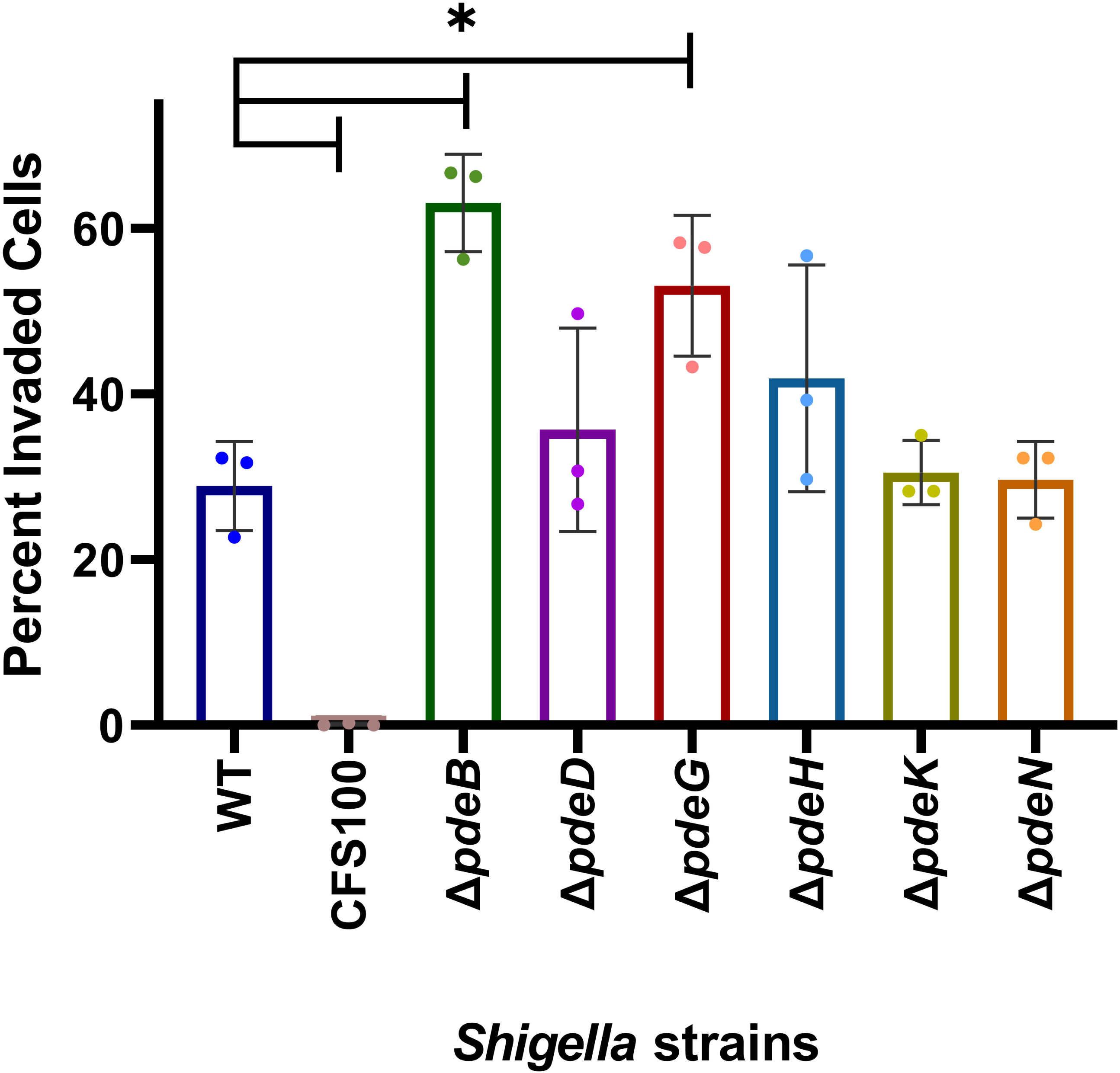
*S.flexneri* Δ*pdeB* and Δ*pdeG* strains exhibited increased host cell invasion. *S.flexneri* strains were used to infect Henle-407 cells and the percentage of invaded cells was determined by Wright-Geimsa staining and microscopy. CFS100 was included as a negative control. Each symbol represents an independent replicate. Bars indicate the mean while the error bars indicate SD. Statistical analysis was performed using One-Way ANOVA with Dunnett’s multiple comparison tests (p < 0.05) to compared to WT. * indicates statistical significance to WT.

### *S.flexneri pdeG, pdeK* and *pdeN* regulate plaque formation

*S. flexneri* DGC mutant strains have a reduced plaque phenotype (35,36). Therefore, we wanted to determine if *Shigella*’s PDEs contribute to *S. flexneri* intercellular spread or intracellular growth. To this end, we performed a plaque assay, using the Henle-407 tissue culture model of infection with gentamycin protection. We infected Henle-407 cells with our 6 *S*. *flexneri* PDE mutant strains and measured plaque size after 48 hours. As negative controls, we included the CFS100 strain lacking the virulence plasmid, and an Δ*icsA* strain that is incapable of cell to cell spread due to its inability to polymerize host actin (56,57).

The *S. flexneri* Δ*pdeG* and Δ*pdeN* mutant strains had a very modest but statistically significant increase in plaque size, while the mutated Δ*pdeK* strain plaque sizes were modestly smaller but statistically significant compared to WT (Fig 5). Our negative controls CFS100 and Δ*icsA* did not form any plaques. The *S. flexneri pdeB*, *pdeD*, *pdeG* and *pdeK* mutants expressing their respective genes showed reductions in plaque size, while *pdeH* and *pdeN* mutants expressing their respective genes showed increased plaque size which were statically significant to WT (Fig S4). We conclude that *Shigella’*s *pdeG*, *pdeK* and *pdeN* regulate the ability to form plaques in Henle-407 monolayers.

**Figure 5.**
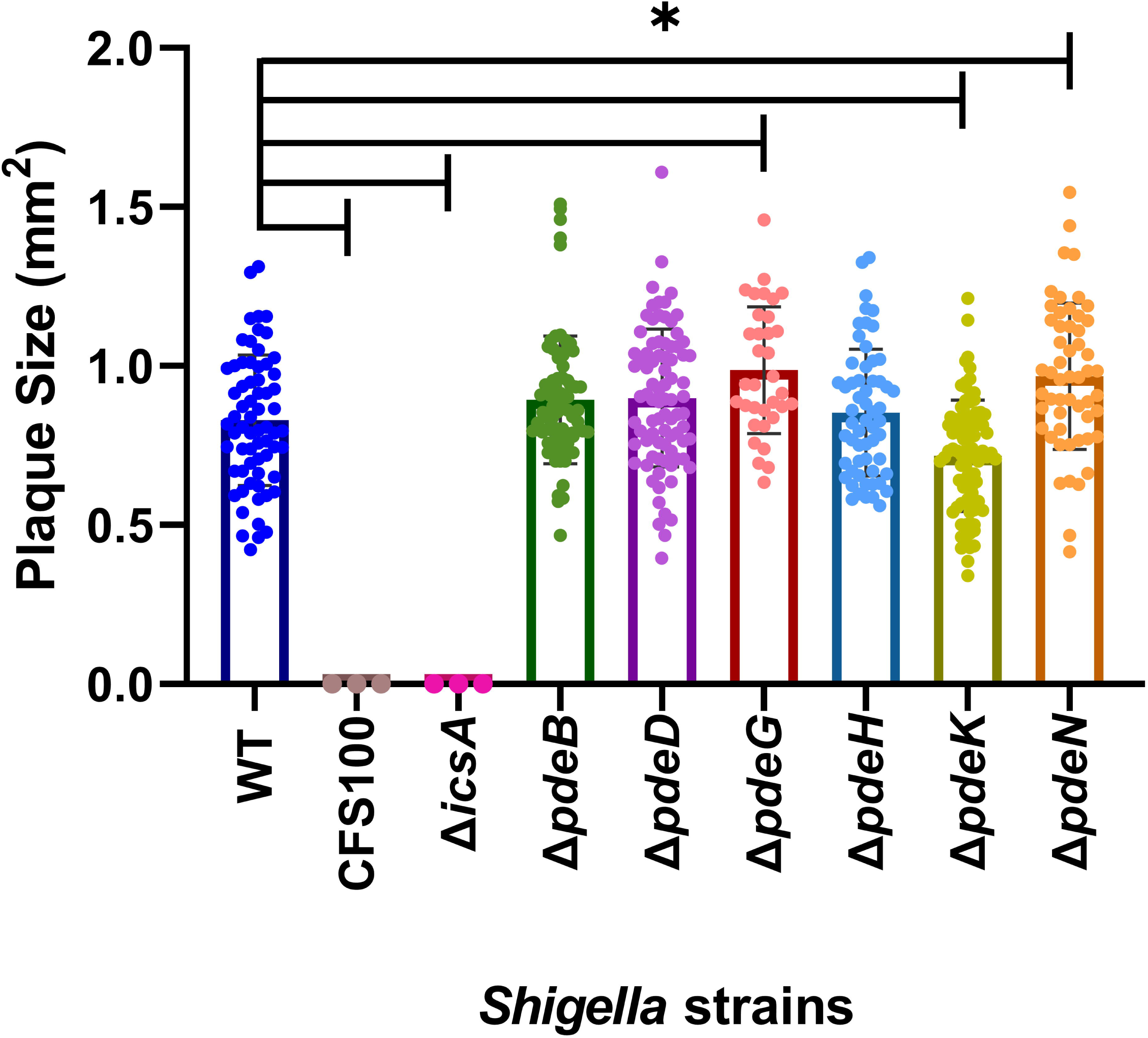
*S.flexneri* Δ*pdeG* and Δ*pdeN* strains exhibited modestly increased plaque size. *S.flexneri* strains were used to infect Henle-407 cells and their plaque size (mm) was quantified. CFS100 and Δi*csA* were included as negative controls. Graph represents a combination of three independent replicates. Each symbol on each respective *S.flexneri* strain represents the size of a plaque. The bar graphs show the mean plaque size while the error bars indicate SD. Statistical analysis was perform using One-Way ANOVA with Dunnett’s multiple comparison tests (p <0.05) to compare WT to CFS100, Δi*csA* and PDE mutant strains. # indicated statistical significance to WT.

### *Shigella* PDEs reduce c-di-GMP levels

Because *S. flexneri* PDE mutants exhibit differences in biofilm formation, invasion, and plaque formation, we postulated that these PDEs are regulating c-di-GMP levels. To determine how *Shigella*’s PDEs impact c-di-GMP levels, we used liquid chromatography tandem mass spectrometry (LC MS-MS) to measure *Shigella* c-di-GMP, which can accurately quantify c-di-GMP with high sensitivity (58,59). Because PDE’s convert c-di-GMP to pGpG, we expected that knocking out *Shigella*’s PDEs will increase c-di-GMP levels within the bacterial cell. This is what we observed, as the *S. flexneri* Δ*pdeB*, Δ*pdeD* and Δ*pdeH* mutant strains had statistically significant increases in c-di-GMP levels (Fig 6A). The strain with the highest c-di-GMP levels was the *S. flexneri* Δ*pdeH* mutant. In *E.coli*, *PdeH* is known to reduce cellular levels of c-di-GMP (60,61), and an *E. coli* Δ*pde*H mutant has been shown to have increased cellular levels of c-di-GMP in *E.coli* (44,61), consistent with our findings.

**Figure 6.**
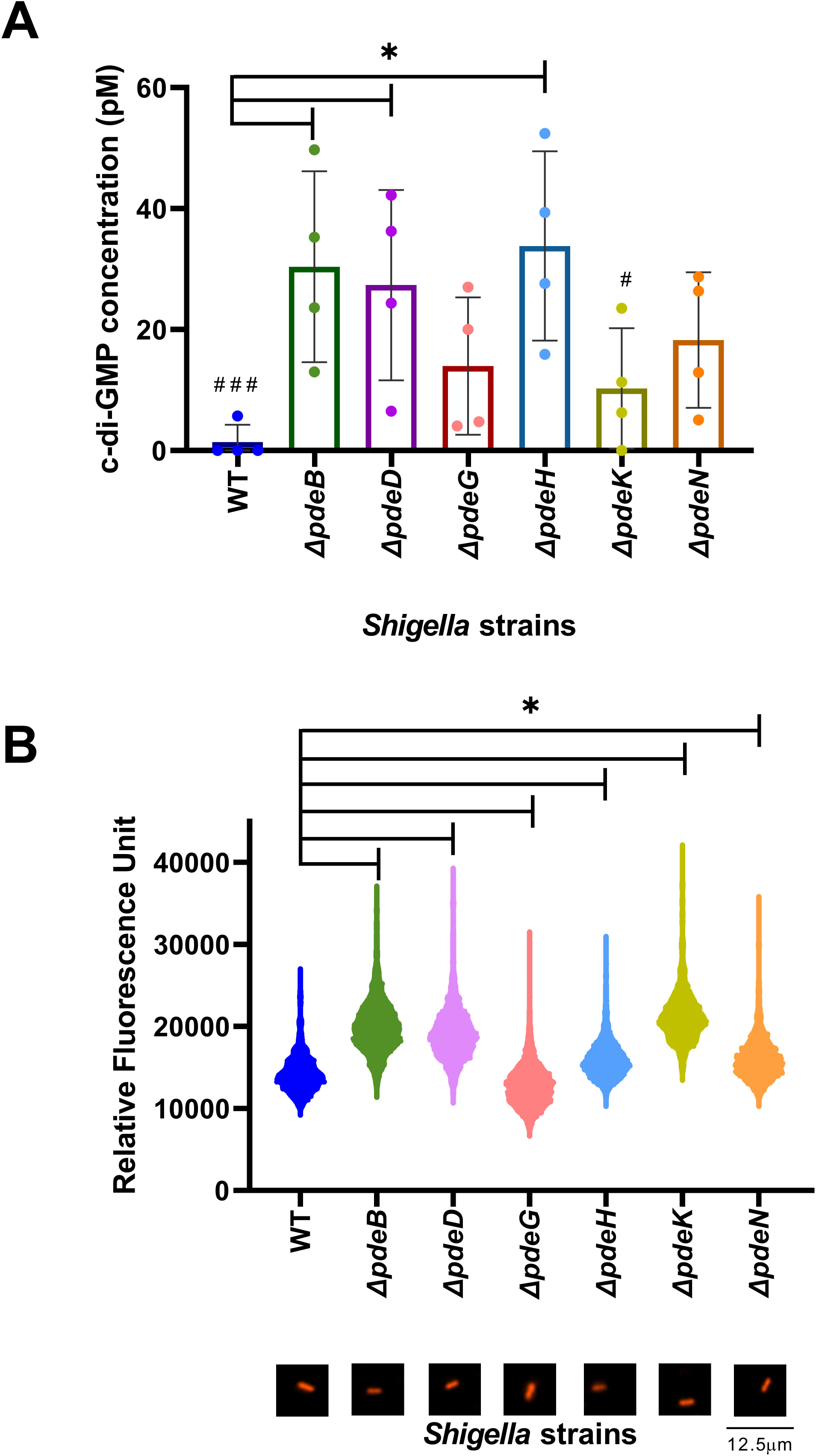
*S.flexneri* mutant PDE strains modulate c-di-GMP levels. (A) LC MS-MS was used to quantify the cellular levels of c-di-GMP within *Shigella* PDE mutant strains. Each symbol represents an independent trial. Statistical analysis performed using One-Way ANOVA with Dunnett’s multiple comparison tests (p < 0.05), compared to WT. The bar graphs show the mean c-di-GMP levels while the error bars indicate SD (B) *Shigella* PDE mutant strains containing a plasmid with a double tandem c-di-GMP riboswitch regulating mScarletI was used to quantify relative c-di-GMP levels by microscopy. This graph represents one out of three independent replicates. Each symbol represents the relative fluorsence unit (RFU) of a single bacterial cell. Micrographs underneath each strain label is an individual bacterial cell. Statistical analysis was performed using a One-Way ANOVA with Dunnett’s multiple comparisons test (p < 0.05) to WT. * indicates statistical significance to WT. # indicates a replicate where c-di-GMP levels were below the detection limit.

We also applied an orthogonal approach to confirm that *Shigella* PDEs regulate c-di-GMP levels, using a biosensor which contains a c-di-GMP double tandem riboswitch (Bc3-4) controlling the translation of the fluorescent reporter protein mScarlet I (62). This approach allows relative c-di-GMP quantitation in live cells, at single-cell resolution. We measured fluorescence of exponentially growing bacteria in LB media supplemented with bile and glucose. We observed that *S. flexneri* Δ*pdeB*, Δ*pdeD*, Δ*pdeH*, Δ*pdeK* and Δ*pdeN* mutant strains exhibited increased fluorescence, indicating increased intracellular levels of c-di-GMP compared to WT (Fig 6B). This data shows that *S. flexneri’s* PDEs are active and play a role in regulating the levels of c-di-GMP within the bacterial cell.

### *Shigella*’s PDEs modulation of c-di-GMP affects cell length

While quantifying changes in our c-di-GMP reporter strains, we observed that some of our PDE mutant strains were consistently biased towards smaller cell lengths. *Shigella*’s bacterial cells are typically between 1-3μM in length, depending on laboratory growth conditions (63). We observed that there were modest decreases in the cell length between WT and all mutant strains with the exception of mutant *ΔpdeG* in the growth conditions used here (Fig 7). This is not the first time that c-di-GMP levels have been linked to cell length; in *Anabaena*, the overexpression of *pde*H from *E.coli* reduces c-di-GMP levels and also reduces cell length and overall cell size (64). We also expressed each of these PDEs in their respective mutant strains. When we compared cell length between each mutant strain and their respective complementation strain, we observed that there were mostly increases in cell length when PDEs were expressed, with the exception of *S.flexneri* Δ*pdeH* and Δ*pdeN* strains and their respective complementation strains, which demonstrated the opposite pattern. *S.flexneri* Δ*pdeG* and Δ*pdeK* + *pdeK* had cell sizes similar to WT (Fig 7). Our data indicates that altering PDE expression can have an impact *S. flexneri* cell length.

**Figure 7.**
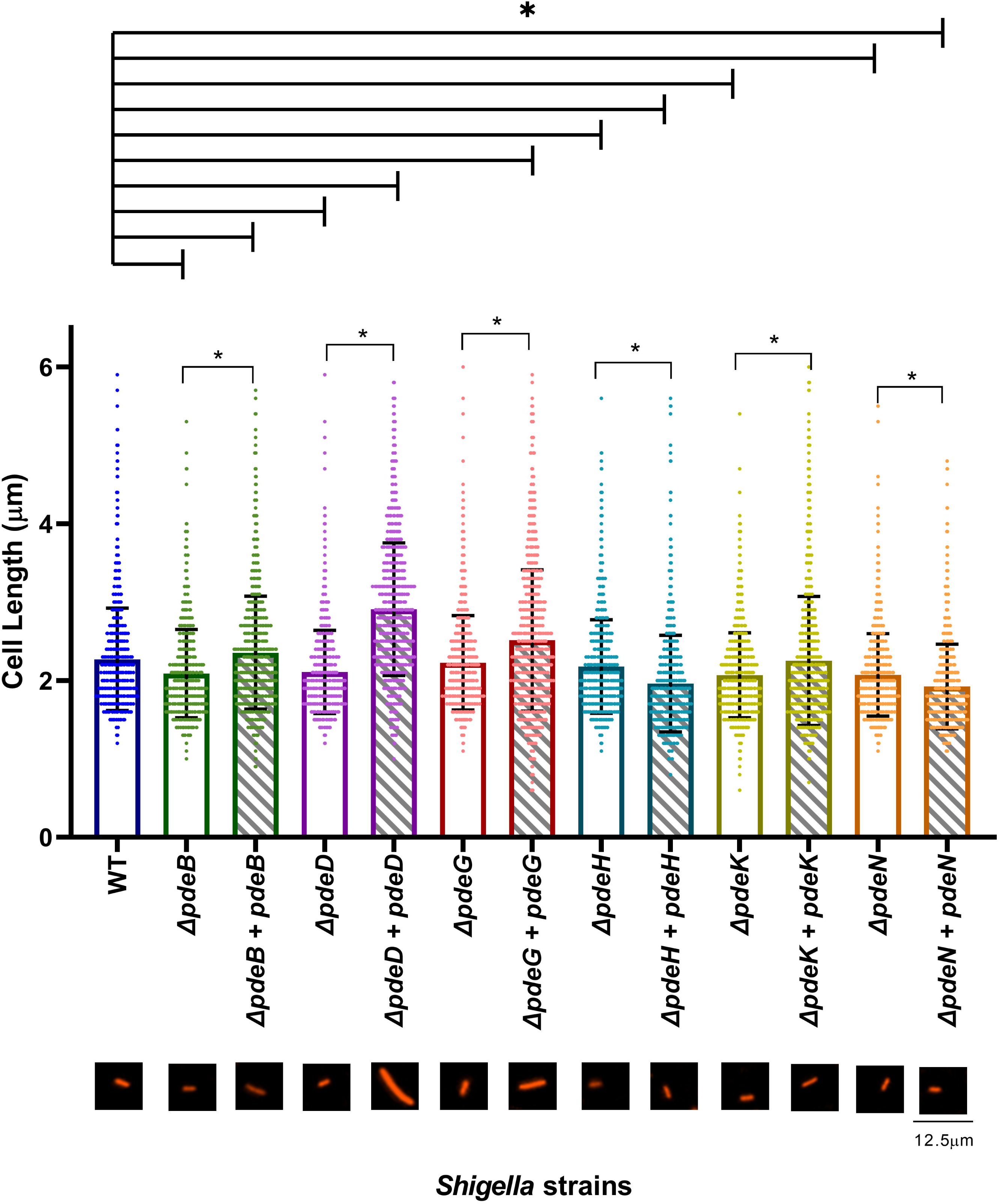
*S.flexneri* PDEs modulate cell length. *S.flexneri* mutant and complementation strains cell length were measured by microscopy from three independent experiments. Bars indicate the mean cell length while the error bars indicate SD. Statistical analysis was performed using a One-way ANOVA with Dunnett’s multiple comparison tests (p <0.05) to compare WT to the mutant and complementation strains. One-way ANOVA with Sidak’s multiple comparison tests (p <0.05) was used to compare each mutant strain to their respective complementation strain.The micrographs underneath each strain label is a representative individual bacterial cell showing cell length. **\*** indicates which strains were statically significant to WT. * indicates which mutant strain was statically significant to its complementation strain.

### *Shigella* Δ*pdeB* differentially expresses genes involved in virulence and lipid metabolism

Because we observed that the *S. flexneri* Δ*pdeB* strain had an increased invasion phenotype (Fig. 4), we performed RNA-seq to identify differentially expressed genes that could contribute to this behaviour. Strains were grown in the same growth conditions used for our invasion assay, and RNA was collected from exponentially growing cells and sequenced using a 300 flow cell kit on the Illumina NextSeq2000 platform resulting in 12 million 2×150bp paired reads per sample (Seqcoast). We determined differential expression using DESeq2 (65). We found that there were over 200 genes differentially expressed in the *S. flexneri* Δ*pdeB* strain relative to the WT strain (Fig 8A and Table S3). We then used Gene Set Enrichment Analysis (GSEA) to identify pathways that were up- or down-regulated. We noted from our GSEA analysis that *Shigella*’s pathogenicity pathway was upregrulated (Fig 8B). From this pathway, *ipgD* and *ipgE* were statisitically significant to WT (Fig 8A and S6). These genes are both encoded on the virulence plasmid. *ipgD* is a T3SS effector protein gene that encodes a phosphatidylinositol 4-phosphatase which is cotranscribed with its chaperone gene *igpE* (66). IpgD reduces host PI4,5P_2_, resulting in ruffling of the host cell membrane that facilitates bacterial invasion (12,24,66,67).

**Figure 8.**
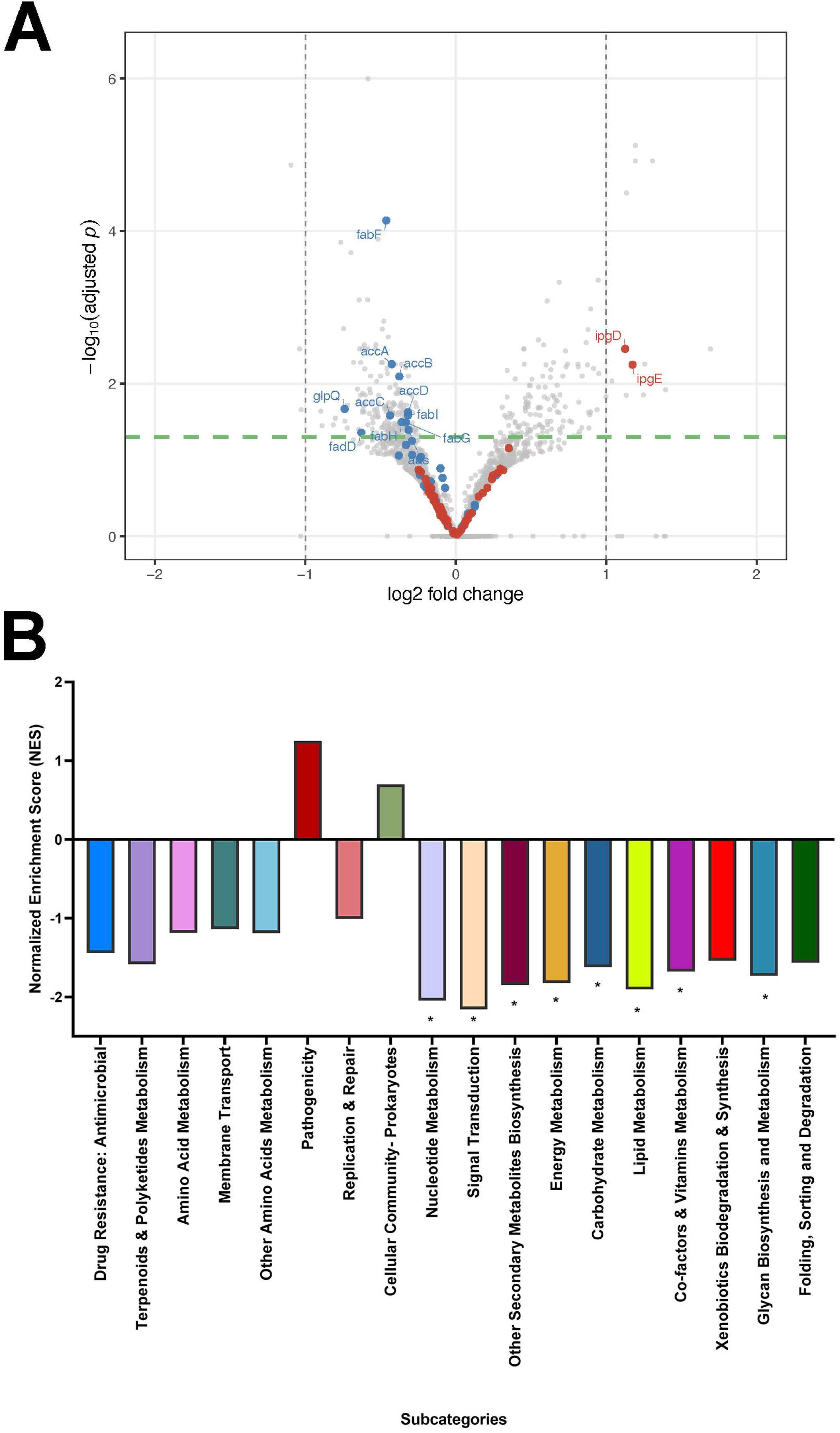
*S.flexneri* Δ*pdeB* differentially regulated over 200 genes compared to WT. (A) Volcano plot showing RNA-seq analysis of differential gene expression between WT and *S.flexneri* Δ*pdeB* strain (high c-di-GMP level) based on their −log_10_ P-value and log_2_ Fold change. All the dots above the threshold line (green) represent genes that were differential expressed compared to WT. A total of 324 genes were differential expressed with 189 genes being downregulated and 135 genes upregulated. The red dots represent all pathogenicity genes and the blue dot represent FA genes. (B) Subcategories from KEGG Pathway of Analysis which were differentially expressed in the *S. flexneri* Δ*pdeB* strain, compared to WT were plotted with their Normalized Enrichment Score. Pathways with a positive value above zero are deemed to be upregulated while those pathways with a negative value below zero are downregulated. * indicates statistical significance to WT. For a detail list of genes that were differentially regulated please refer to supplemental Table S3.

We also noted other KEGG pathway subcategories were down-regulated in the *S. flexneri* Δ*pdeB* strain, including nucleotide metabolism, signal transduction, secondary metabolites, energy metabolism, lipid metabolism, cofactor metabolism, xenobiotic degradation, and glycan biosynthesis (Fig 8B). We focused on lipid metabolism, where the fatty acid (FA) synthesis genes *Acc*(*ABCD*), *fab*(*FGHI*), as well as FA degradation gene *fadD* (68) were all expressed at significantly lower levels than the WT strain. Owing to the fact that many of the FA metabolism genes that are repressed in the *S. flexneri* Δ*pdeB* strain are involved in both biosynthesis and degradation of FA, it is difficult to predict the net effect of these changes in gene expression on bacterial lipids. *fadD* was downregulated in both FA synthesis and degradation in our RNA seq data. Knocking out *fad*D in *E.coli* has been shown to increase cytosolic FAs (69). FA metabolism has been previously linked to *S. flexneri* virulence, by directly binding and inhibiting the activity of the master transcriptional regulator *VirF* (70). We created a *S. flexneri* Δ*fad*D strain and found that it had an attenuated invasion phenotype and a miniscule decrease in it’s ability to spread intercellularly (Fig 9). We also noted that this strain exhibited a pink colony phenotype on TSBA + Congo red plates (Fig S7), an indicator that the *Shigella* T3SS is not being expressed normally (53). We confirmed the presence of an intact virulence plasmid in our Δ*fad*D strain by PCR and gel electrophoresis (data not shown).

**Figure 9.**
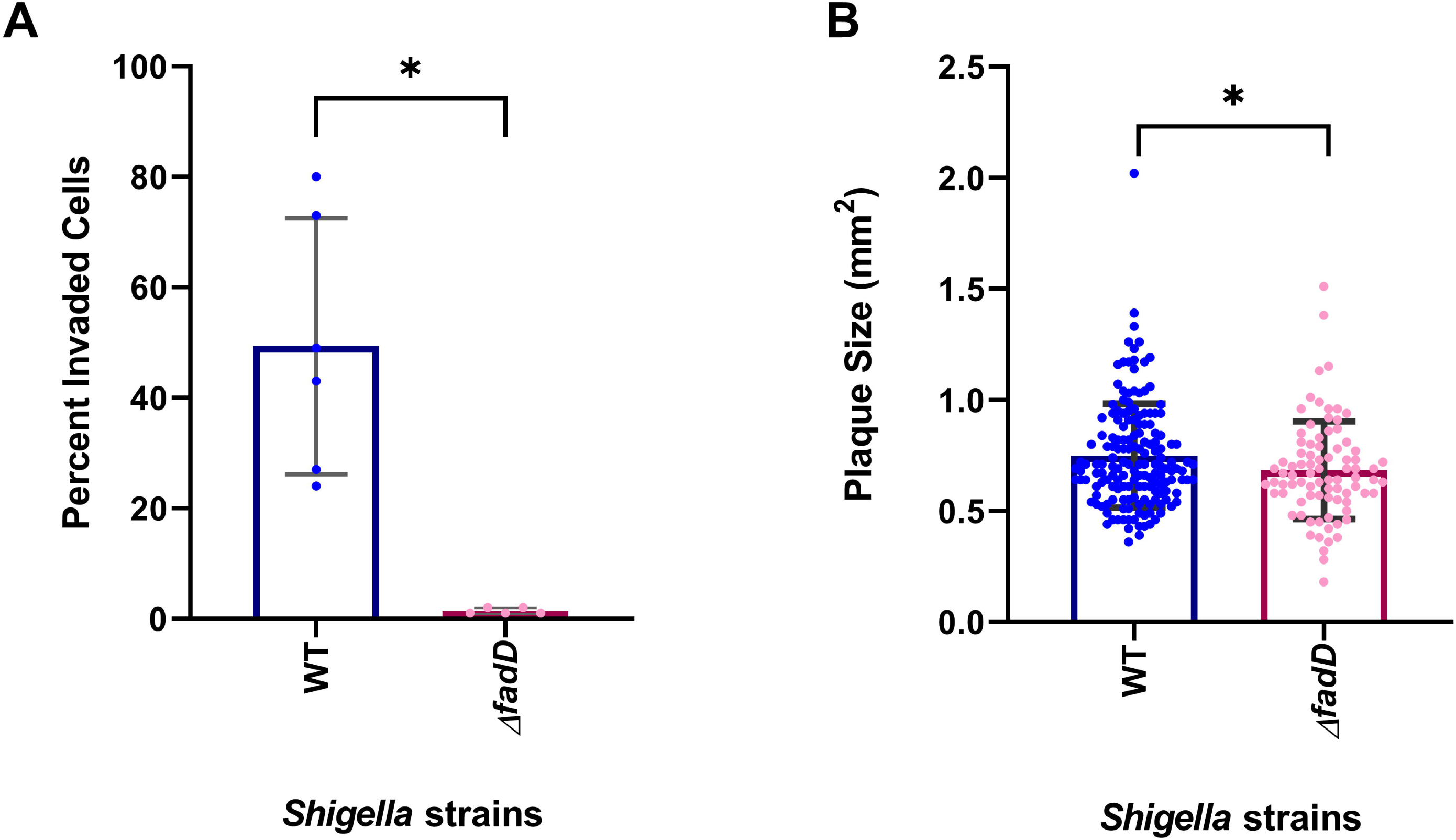
Deletion of *fadD* reduces *S. flexneri* invasion and plaque formation in Henle-407 cells. (A) Quanitification of invasion frequency in Henle-407 cells infected with WT and the *Shigella* Δ*fadD* mutant strains. The *S.flexneri* Δ*fadD* strain exhibited a significant decrease in the percentage of invaded host cells compared to WT. (B) Plaque formation assay in Henle-407 cells infected with WT and the *Shigella* Δ*fadD* strains. Individual data points correspond to plaque sizes for each strain. The *Shigella* Δ*fadD* mutant produced fewer plaques than WT and displayed a modest but statistically significant reduction in plaque size. Bars represent mean ± statistical difference from three independent biological replicates. Statistical analysis was performed used unpaired T-test with Welch’s correction (p < 0.05).

We also wanted to determine the net effect that c-di-GMP mediated changes in FA gene expression have on *S. flexneri* lipid molecules. We determined the FA profile of *S. flexneri* WT, Δ*pdeB,* Δ*pdeB+B* and Δ*fadD* strains using gas chromatograph mass spectrometry (GC-MS). FAs were extracted from *S. flexneri* WT, Δ*pdeB,* Δ*pdeB+B* and Δ*fadD* strains and converted to fatty acid methyl esters as previously described (71). We supplemented every sample with heptadecanoic acid during the FA extraction as an internal spike-in control. We observed our internal control as well as C14, C16:1, C16, and C17:1 (Fig.10 & Table S2). We noted that the *S. flexneri* Δ*pdeB* strain had reduced C18 FAs compared to WT. When *pdeB* was expressed in the *S. flexneri* Δ*pdeB* mutant strain (Δ*pdeB+B*), we observed no statistical significance to WT. Interestingly, we did not observe an abundance of FAs in our Δ*fadD* strain. As such we decided to grow our four *S.flexneri* strain overnight to determine if this allow for greater cytosolic FAs accumulation as seen in *E.coli ΔfadD* strain. We noticed an additional two FAs C15 and C17 being produced in addition to C14, C16:1, C16, C18:1 and C18 (Fig S8). However, *S.flexneri ΔfadD* strain had an increase in C15 and C16 FAs when compared to WT (Fig.S8 ii and iv). This data shows that c-di-GMP is playing a role in modulating FA gene expression and consequently lipid molecules by a mechanism we do not know.

**Figure 10.**
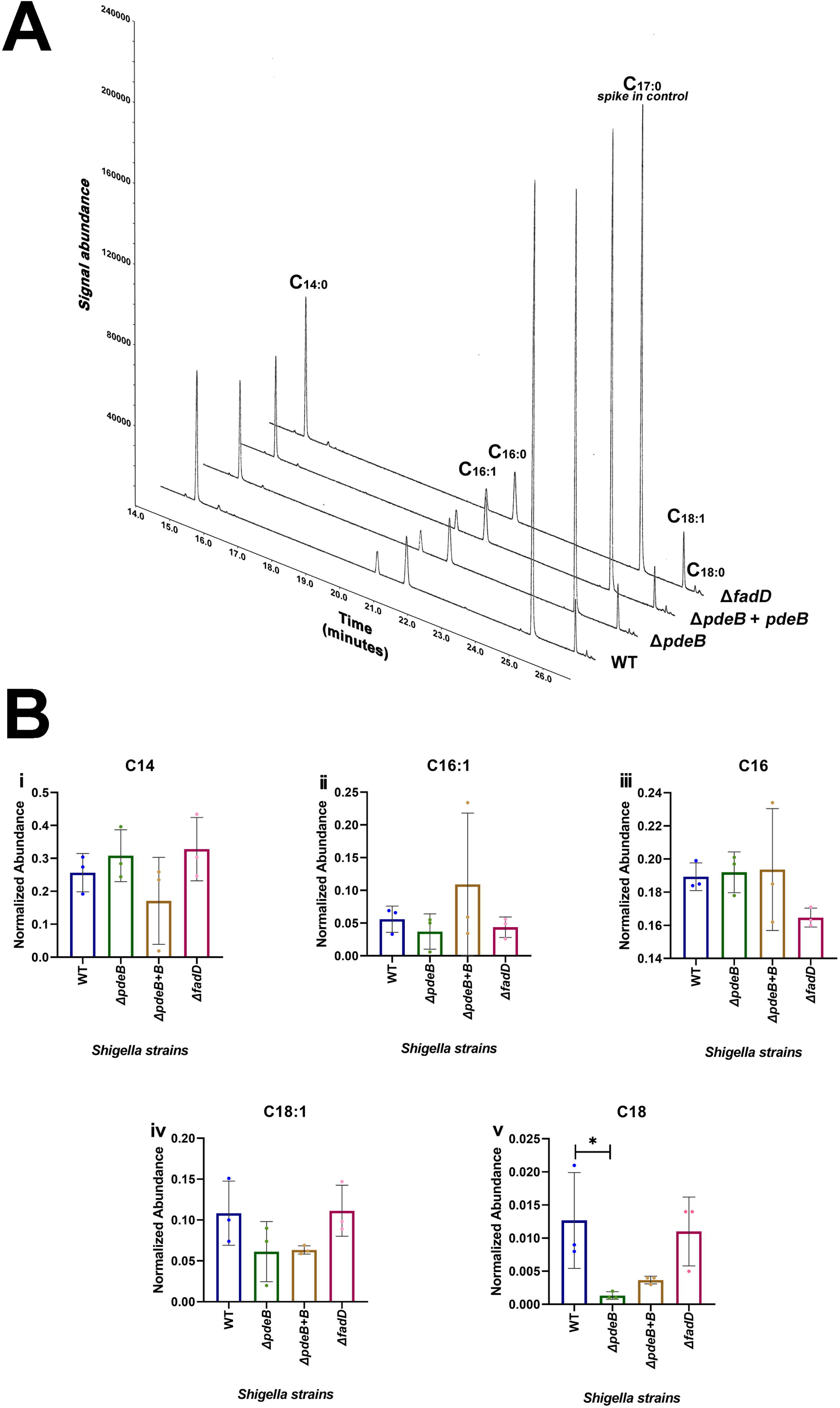
*S. flexneri pdeB* modulates lipid molecules. (A) The GC-MS spectra showing peaks for four FA methyl esters: C15, C17:1, C17, C19:1 and C19 and heptadecanoic acid (C18) as our internal control. (B) For each FAs observed, normalised abundance was calculated and statistical significance to WT was performed using One-Way ANOVA with Dunnett’s multiple comparisons test (p < 0.05). * indicates statistical significance to WT.

**Figure 11.**
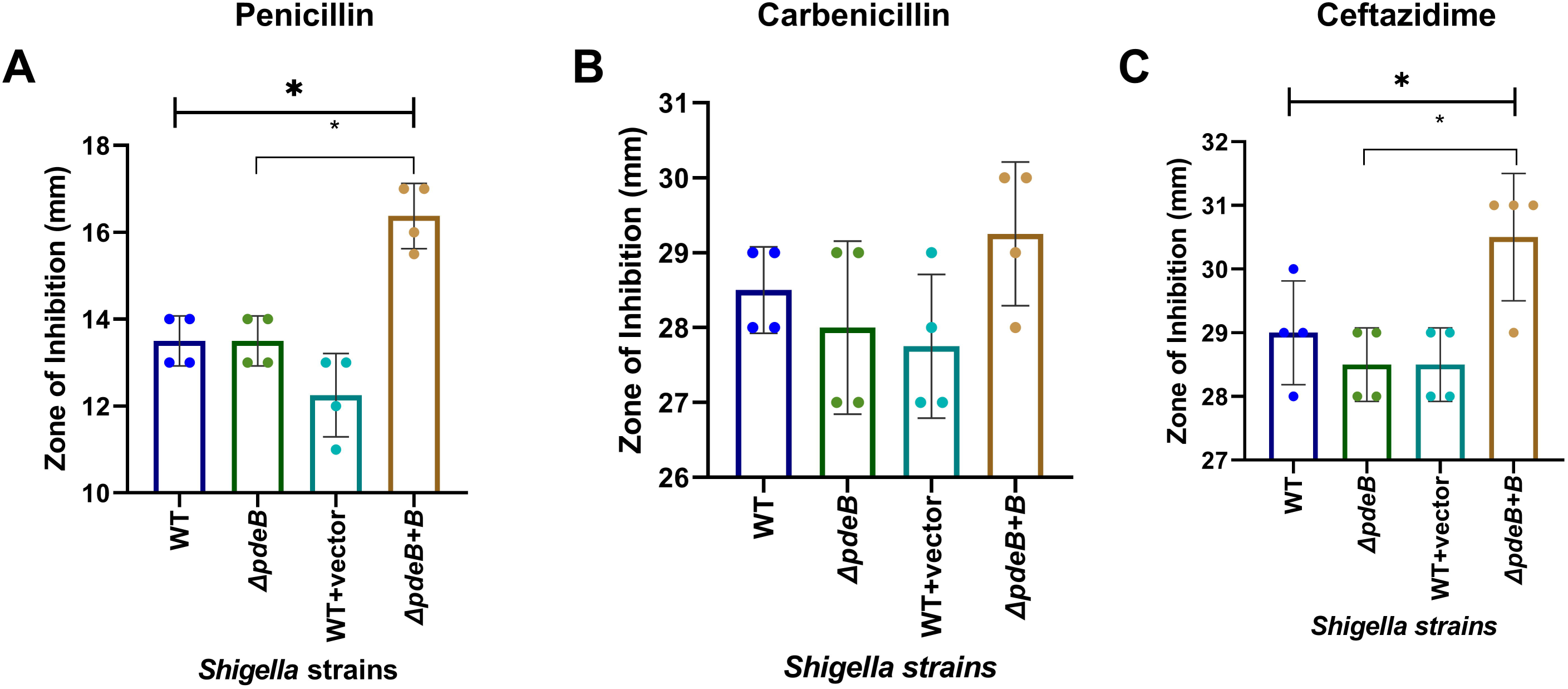
*pdeB* expression increases *S. flexneri* antibiotic sensitivity. Antibiotic sensitivity of WT, *S.flexneri* Δ*pdeB,* WT + vector and *S.flexneri* Δ*pdeB+B* to (A) penicillin, (B) Carbenicillin and (C) Cetfazidime was performed using the Kirby Bauer Disk Diffusion Assay. Zones of inhibition was measured for four independent trials. Bars indicate the mean zone of inhibition, while the error bars indicate SD. Statistical analysis was performed using a One-way ANOVA with Dunnett’s multiple comparison tests (p <0.05) to compare WT to *S.flexneri* Δ*pdeB,* WT + empty vector and *S.flexneri* Δ*pdeB+B* strains. Statistical analysis comparing *S.flexneri* Δ*pdeB* strain to *S.flexneri* Δ*pdeB+B* strain was performed used unpaired T-test with Welch’s correction (p < 0.05). **\*** shows stastical significance to WT. * shows statistical significance to *S.flexneri* Δ*pdeB* strain.

### *pdeB* expression increases *Shigella*’s antibiotic sensitivity

Transcriptional regulation of lipid synthesis can impact the structure of the bacterial lipid membrane and have a significant effect on antibiotic resistance (72,73). Because we observed that modulating c-di-GMP can affect cell size, FA synthesis and in-turn lipid molecules, we wanted to determine if these changes are associated with changes to *Shigella*’s antibiotic sensitivity. We performed a Kirby Bauer disk diffusion assay using ß-lactams (carbenicillin, penicillin and ceftazidime) on WT, *S.flexneri* Δ*pdeB* and *S.flexneri* Δ*pdeB +B* strains. We included WT containing an empty vector (WT + vector) as a control for *S.flexneri* Δ*pdeB+B* strain *to* ensure that the Kan cassette on pVES143 is not affecting antimicrobial sensitivity during the experiment. From our experiment, *S.flexneri* Δ*pdeB*, WT + vector strain showed similar zones of inhibition compared to WT. We saw that when we expressed *pdeB* in its mutant background, there was an increase in the zone of inhibition with penicillin and ceftazidime which was statistically significant to WT. This data shows that c-di-GMP can have an impact on *Shigella*’s sensitivity to antibiotics.

## Discussion

C-di-GMP is a global regulator that allows bacteria to rapidly respond to different environments, facilitating lifestyle changes (47,74,75). We previously found that *S. flexneri*’s four intact DGC’s that are homologous to *E. coli* regulate its biofilm, acid resistance and virulence phenotypes, while modulating the levels of c-di-GMP within the bacterial cell (35,36). Biofilms are associated with elevated levels of c-di-GMP in many different bacteria (76). *E. coli* biofilms are often composed of curli, polysaccharides and cellulose, all of which are induced by high c-di-GMP levels (76,77). In *Shigella* however, many of these genes involved in forming biofilm in *E.coli* are disrupted due to deletions, insertations or frameshift mutations (78–80). Despite this, c-di-GMP can still induce biofilm formation in *Shigella* (36). Here we observed that our S*. flexneri* PDE mutant strains produced higher biofilm levels, while the expression of these PDE’s in their respective mutant background strains had the opposite effect (Fig. 3, Fig. S1). This data is consistent with S*. flexneri* ΔDGC mutants exhibiting reduced biofilm formation (36). Similarly, an *E.coli* Δ*pdeH* strain has higher levels of c-di-GMP and biofilm formation, (44,81) and *E. coli* PdeK reduces c-di-GMP produced by *DgcC* and has a negative impact on microcolony biofilm (42). We conclude that *S. flexneri* PDEs contribute to the negative regulation of biofilm formation.

Different pathogens use c-di-GMP to regulate virulence phenotypes in different ways (58,82). In bacteria like *Salmonella, Klebsiella pneumoniae* and *V. cholera,* low c-di-GMP levels favors invasion while high c-di-GMP levels result in an attenuated phenotype (83–86). Overexpressing the DGC VCA0956 in *S. flexneri* increases c-di-GMP beyond normal physiological conditions and results in a reduced invasion and plaque phenotype; however, mutating *S. flexneri dgcF* reduces c-di-GMP levels, and also reduces invasion and plaque formation (35,36). To determine the impact of PDEs on *Shigella*’s virulence, we used host cell invasion and intercellular spread as markers for virulence. Consistent with *Shigella*’s DGCs contributing to virulence regulation, specific PDEs are also regulating *Shigella* virulence phenotypes. *S. flexneri* Δ*pdeB* strain exhibits increased invasion (Fig. 4), while *S.flexneri* Δ*pdeG* strain had increased invasion and cell to cell spread (Fig 4 and Fig 5). Transcriptome analysis revealed that the *S. flexneri* Δ*pdeB* strain had an increase in gene expression of the effector protein *ipgD* and its chaperone *ipgE* (Fig.S6), which could possibly contribute to this observed behaviour.

We also observed that c-di-GMP represses the expression of genes associated with *Shigella*’s lipid metabolism (Fig 8). This is not the first time c-di-GMP has been implicated in regulating FA gene expression. In *Mycobacteria smegmatis*, c-di-GMP binds to a nucleoid associated protein *Lsr2* (87), and this binding results in the expression of *16ad*, which is essential for synthesing full length mycolic acids (87,88). Consistent with *pdeB* differentially regulating FA gene expression, we observed decreases in a C18 (stearic acid) FA in our *S.flexneri* Δ*pdeB* strain, while *S.flexneri* Δ*pdeB+B* strain resulted in this FA not being statistically significant to WT (Fig.10v). FAs such as palmitoleic acid, oleic acid and stearic acid can bind to VirF, a virulence regulator in *Shigella* (89). This binding resulted in reduced transcription of *virB* which promotes the transcription of genes involved in host cell invasion such as the T3SS, *mixE* and other virulence effectors (90–92); this ultimately reduces *Shigella*’s invasion capability (89). It is possible that lower levels of endogenous C18 FA observed in the *S. flexneri* Δ*pdeB* strain is contributing to the upregulation of virulence genes associated with *Shigella*’s virulence.

In *E.coli*, a Δ*fadD* mutation increases cytosolic FA (68). Our *S.flexneri* Δ*fadD* strain did not show significant acculmulation of FAs as we expected when grown to exponential phase as we would when performing our invasion assay. FA profiles however, can vary among strains (93), and accumulation of FAs are typically seen during late stationary phase (71). When we performed FA extraction during stationary phase, we noted an increase in C15 and C16 FAs in the *S.flexneri* Δ*fadD* strain compared to WT (Fig S8 ii and iv). We also noted reduced invasion and plaque phentotypes for our *S.flexneri* Δ*fadD* strain, consistent with fatty acids altering *S. flexneri* virulence phenotypes (89)(Fig 9). Disruption to *fadD* have resulted in repression of virulence genes such as *ctxAB* and *tcpA* in *V.cholerae* and *hilA* in *Salmonella enterica* (94,95).

Mutation of *S. flexneri* PDEs resulted in increased c-di-GMP levels compared to the WT strain. We noted that altering *S. flexneri* PDE expression resulted in differences in cell length. Bacterial cell size is dependent upon cell growth and division (96). In pathogens such *Caulobacter crescentus* and *Anabaena*, c-di-GMP plays a role in cell cycle progression and maintaining cell size (64,97). Most of our *S. flexneri* PDE mutant strains had modest but consistent reductions in cell length, and PDE expression increased cell length in the same modest and consistent manner. The *S. flexneri pdeD* expression strain had the most notable cell length difference, that was almost 25% longer than WT. This is not the first example of a c-di-GMP turnover enzyme regulating cell length (48,98). *E.coli’*s *dgcN* has been shown to modulate intracellular levels of c-di-GMP, and *Salmonella* cells expressing *dgcN* had modest increase in cell length and no mid-cell invagination, indicating that this DGC could be restricting Z ring constriction and synthesis of septal peptidoglycan (48). Further studies are necessary to understand the mechanism by which *Shigella*’s PDEs are modulating c-di-GMP levels to affect cell length.

Our *S.flexneri* Δ*pdeB* strain showed decreased *fab* gene expression and lipid molecules. Lipid content of bacterial membranes can impact antimicrobial’s efficacy (99). Additionally, regulating c-di-GMP levels in the bacterial cell can play a role in an antimicrobial resistance. High c-di-GMP levels have been associated with antibiotic resistance while lower levels of c-di-GMP results in greater antimicrobial susceptibility (100,101). Our data is congruent with this as our *S.flexneri* ΔpdeB strain which increased c-di-GMP levels had reduced susceptibility to the ß-lactams we tested. When we expressed *PdeB* which negatively regulates c-di-GMP we saw increased susceptibility to penicillin and ceftazidime.

## Methodology

### *S.flexneri* growth conditions and media

Tryptic soy broth (TSB) plates with 0.01% Congo red was used to culture all *S.flexneri* strains used in this study. Strains were grown at 37°C and red colonies were selected to ensure the virulence plasmid was present (55,102). Overnight cultures were grown in Luria Betani (LB) broth shaken at 30°C.

For strains carrying pVES143-derivative plasmids, media was supplemented with 50µg/mL Kanamycin (Kan) and 2mM Isopropyl-ß-D-thiogalactopyranoside (IPTG) for plasmid maintenance and induction, respectively. Unless otherwise stated, liquid media used for experimental assays and RNA extraction were supplemented with 0.05% (wt/vol) sodium deoxycholate (DOC) (Sigma-Alridch).

Henle-407 cells (ATTC, CCL6) were used as our tissue culture model for the virulence assays. Henle-407 cells were routinely maintained with Minimal Essential Media (MEM, Gibco Catalog No. 61100087) supplemented with 10% (wt/vol) BactoTryptone Phosphate broth (Difco), 10% fetal bovine serum (Corning Catalog No. 35-053-CM), 2mM glutamine (Gibco GlutaMAX Supplement Catalog No. 35-050-061), 0.55**%** (wt/vol) Sodium Bicarbonate, and MEM non-essential amino acids (Gibco Catalog No. 11-140-050).

### *S.flexneri* strains, mutants and plasmids generated

*S.flexneri* 2457T was the parental strain used to create all PDE mutants in this study (Table S4), using the one step inactivation of chromosomal genes as described by Wanner and Datsenko (46). PCR was used to amplify the chloroamphenicol (Cam) cassette of pkD3 with overhangs homologous to our genes of interest (Table S5). Gene knockouts were verified by Sanger sequencing.

The *S. flexneri* Δ*fadD* mutant strain was generated by P1 bacteriophage-mediated transduction using the *E. coli* Δ*fadD* strain from the Keio collection (Table S4) (103,104). After P1 transduction, the mutant strain was selected on TSBA-CR containing 50 µg/mL kanamycin. The mutation was then validated by PCR amplification (Table S5) and sequencing (Azenta). The virulence plasmid in the *Shigella* Δ*fadD* mutant was confirmed by PCR amplification using *phoN1* primers (Table S5), followed by agarose gel electrophoresis of the PCR product (Figure S7).

For our complementation strains (Table S4), we cloned each PDE into the pVES143 plasmid using the NEBuilder HiFi DNAssembly kit (New England Biolabs Catalog No. E2621L), using primers listed in Table S5. We then transformed these plasmids into their respective mutant strains by chemically induced competence. Strains were selected for on Kan selection plates and confirmed by whole plasmid sequencing.

### Growth Curve Assay

Overnight cultures were sub-cultured 1:50 in M63 medium supplemented with 10mM glucose, nicotinic acid and 10mM magnesium sulfate (Mg_2_SO_4_). 150µL of bacterial suspension was added to a 96 well plate. Plates were incubated at 37°C with shaking and the OD_600_ was read every hour for over 24 hours.

### Biofilm Assay

Biofilm micro-titre assay was performed as described by Nickerson and Faherty (50), with minor alterations. Overnight cultures were diluted 1:50 in LB containing 0.4% sodium deoxycholate (DOC) (Sigma) and 2% glucose. For the strains containing pVES143, the media was supplemented with Kan and IPTG. Using a 96-well tissue culture treated plate (Fisherbrand Catalog No. FB012931), 130μL of each strain was dispensed in sextuplicates and incubated statically at 37 °C for ∼18 hours. OD_600_ were taken for each sample. The media was decanted and the biofilm formed was washed with PBS (Gibco Catalog No. 10010023) and allowed to dry. 0.5% crystal violet (CV) was used to stain the biofilm which was then solublized in 95% ethanol at 4°C for 30 minutes. Due to high biofilm formation, the solubilized biofilm was diluted 1:10 and the OD_540_ was recorded for each strain. Bioflim assays were repeated three independent times.

### Invasion Assay

This assay was carried out according to Koestler et al. with some modifications (53). Overnight cultures were diluted 1:100 in LB media with 0.05% DOC and grown until mid-log phase at 37°C shaken. A volume of 2 x 10^9^ CFU/mL was centrifuged and resuspended in PBS. Henle 407 cells grown in a 6 well plate to ∼20% confluency was infected with 100μL of cell suspension. Plates were spun for 10 mins at 1000 RPM. Cells were then incubated for 1 hour at 37°C in 5% CO_2_. After incubation cells were washed 4 times in PBS. The cells were then incubated in MEM supplemented with gentamycin for 1 hour. The cells were then washed 4 times and stained with Wright-Giemsa stain. Each strain’s invasion percentage was quantified by microscopy. A total of 300 Henle cells were counted and their invaded positivity determined. Positively invaded cells were counted as any Henle cell with 3 or more bacterial cells. Invasion percentage was then calculated based on the number of positively invaded cell per strain divided by 300. Invasion assays were repeated three independent times.

### Plaque Assay

The plaque assay conducted according to Koestler et al. with some modifications (53). Overnight cultures were diluted 1:100 in LB media with 0.05% DOC and grown until mid-log phase at 37°C shaken. A volume of 5 x 10^4^ CFU/mL was centrifuged and resuspended in PBS. Henle 407 cells grown in a 6 well plate to ∼80% confluency was infected with 100μL of cell suspension. Plates were spun for 10 mins at 1000 RMP. Cells were then incubated for 1 hour at 37°C in 5% CO_2_. After incubation cells were washed 4 times in PBS. The cells were incubated in MEM supplemented with gentamycin and glucose for 48 hours. The cells were then washed 4 times and stained with 0.5% CV. Plaque sizes were imaged using BioRad Gel Doc Imager and quantified using Image J. Plaque assays were repeated three independent times.

### Quantification of c-di-GMP

For LC-MS quantitation of c-di-GMP, overnight cultures were sub-cultured 1:100 in LB media containing 0.05% DOC and 0.1% (vol/vol) glucose until an OD_600_ between 0.4- 0.7 was attained. The cells were pelleted at 4°C for 10 mins at 7000rpm. The pellet was washed once in 1x PBS. The cells were resuspended in 700µL cold extraction buffer comprised of methanol, acetonitrile, water (40:40:20) and 0.1N formic acid. Cells were incubated at −20°C for 30 minutes. The cells were centrifuged at 13,000rpm for 10 minutes at 4°C. The supernatant was retained and dried in a vaccum concentrator. The dried pellet was then resuspended in 50µL of water. C-di-GMP was detected using Waters SYNAPT XS.

For quantitation of c-di-GMP by fluorescence microscopy, overnight cultures were sub-cultured 1:100 in LB supplemented with 0.05% DOC and 50µg/mL of Ampicillin to maintain pRO010 derrived plasmid for ∼2.5 hours. Cultures were then diluted 1:100 in M63 medium supplemented with 10mM glucose, nicotinic acid, 0.05% DOC, 10mM magnesium sulfate (Mg_2_SO_4_) and 50µg/mL of Ampicillin. 100µL of bacterial suspension was added to a 96 well plate (Greiner Catalog No. 655892). The plate was spun at 1000 rpm for 10 minutes. C-di-GMP was then quantified using microscopy exposure settings: illumination intensity 10, integration time 700 and camera gain 20.

### RNA Extraction

Overnight cultures were sub-cultured in a 1:100 dilution in LB till mid log OD_600_. Bacterial cells were then harvested and extracted using Pure Link RNA Mini kit (Cat no. 12183020). RNA quality was determine using Agilent 4150 Tape Station and reagents (RNA Screen Tape Catalog No. 5067-5576, RNA ScreenTape Sample Buffer Catalogy No. 5067-5577). All samples had a RIM value of 8.5 and above. Samples were sent to SeqCoast Genomics for RNA sequencing. Sequences were deposited in the NCBI Gene Expression Omnibus in the study GSE330893 (sample accession numbers GSM9735135 - GSM9735140).

### RNA-seq data processing and differential expression analysis

Paired-end RNA-seq reads were quality-filtered and Illumina adapters were trimmed with BBduk (BBMap v39.01) with default parameters for adapter removal and quality trimming (105). Trimmed reads were aligned to the *Shigella flexneri* 2457T reference genome using BWA-MEM (v0.7.17) (106,107). Resulting alignments were sorted and indexed using SAMtools (v1.18) (108).

Gene-level read counts were generated using featureCounts (Subread v2.1.1) (109). Counting was performed at the coding sequence (CDS) level via annotation derived from the reference genome sequence. Specifically, read pairs were counted using the --countReadPairs option, with filters requiring both mates to be aligned (-B) excluding chimeric fragments (-C). Strand specificity was used for sequencing and the reverse-stranded configuration (-s 2) was used for all downstream analyses.

Lowly expressed genes were filtered prior to differential expression (DE) analysis by requiring ≥ 10 counts in at least three samples. Pairwise DE analyses were performed using DESeq2 (65) (Bioconductor (110)), by condition (e.g. WT vs deltaB). Log_2_ fold changes (FC) were shrunk using the apeglm option in DESeq2 to improve effect size estimation. Genes with an adjusted p-value (Benjamini–Hochberg correction) of ≤ 0.05 were considered significantly differentially expressed to correct for the false discovery rate of false positives. Visualization of DE results as volcano plots were generated using the EnhancedVolcano R package (111).

### Functional enrichment analysis

Functional enrichment analyses were performed using KEGG pathway annotations and the **clusterProfiler** R package (112,113). Gene identifiers from DE results were mapped to KEGG identifiers derived from the reference genome annotation. Gene set enrichment analysis (GSEA) was performed using all genes tested in the DE analysis that were successfully mapped to KEGG identifiers, ranked by log_2_ FC. Enrichment scores were summarized as normalized enrichment scores (NES), and significance was based on adjusted p-values. KEGG pathways were analyzed hierarchically at the levels of category, subcategory, and individual pathway. Results were visualized using bar plots of enrichment significance (−log_10_ adjusted p-value) and gene counts to summarize pathway enrichment across contrasts.

### Fatty Acid Extraction and Analysis

FA extraction was performed according to Politz et al with some modiifications (71). Briefly, Overnight cultures were diluted 1:100 in LB supplemented with 0.05% DOC and grown until mid-log and stationary phase at 37°C shaken. 5µL of heptadecanoic acid (ThermoFisher Scientific Catalog No. 506-12-7) along with 100µL of glacial acetic acid (Fisher Scientific Catalog No. A38-212) was added to bacterial cultures. 5mLs of a mixture of chloroform and methanol [1:1] was then added to the mixture above. The cultures were centrifuged at 1000 x g for 10 mins at 18°C. The aqueous and interface layers were aspirated and the chloroform layer was lyophilized in a speed vac. The anhydrous sample was dissolved in 500µL of 1.25M HCL-methanol (Sigma Aldrich Cat no. 17935-50ML) and incubated at 75°C for 1 hour. 500µL of hexane (Sigma Aldrich Cat no.650552-4L) was added to each sample along with 5mLs of sodium bicarbonate (Fisher Scientific Catalog No. S233-500). The mixture was vortex and centrifuged. The hexane layer containing the FA was retained. 500µL of hexane was added to the tubes which was vortexed, centrifuged and the hexane layer was combined with the hexane layer retained earlier.

GC analysis of the FA extracts was adapted from Ren et al. using Algilent 7683 Series Injector and 5973 Network Mass Selective Detector and Agilent 19091J-433 30m x 250µm x 0.25µm capillary (114). The oven temperature gradient started at 70°C for 2 minutes which was then ramped to 170°C at a rate of 11°C/min and held for 2 minutes. A further temperature increase to 175°C at a rate of 0.8°C/min held for 4 minutes followed by a final temperature increase to 220°C at a rate of 20°C/min held for 2.5 minutes, for a total run time of 28.09 minutes.

### Antibiotic Sensitivity Assay

Antibiotic sensitivity was determined using the Kirby-Bauer Disk Diffusion Susceptibility Test (115). Briefly, overnight cultures were dulited 1:100 in LB supplemented with 0.05% DOC and grown until mid-log phase at 37°C shaken. Cultures were adjusted to give a concentration of 1 x 10^8^ CFU/mL cells, Strains were inoculated onto the surface of Muller Hinton agar (BD Difco Muller Hinton Agar Catalog No. SKU:225250) plates. The plates were allowed to dry. 10µg of Penicillin (Sensi-Disc Penicillin Catalog No. 230918), 100µg Carbenicillin (Sensi-Disc Carbenicillin Catalog No. 231555) and 30µg Cetfazidime (Seni-Disc Ceftazidime Catalog No. 231632) disks were placed on to the plates. Plates were incubated at 37°C for ∼18 hours. Zones on inhibition was measured in milimeters.

### Statical Analysis

Statical analysis was performed using GraphPad Prism. A p value of <0.05 was considered statistically significant.

## Acknowledgements

We would like to thank Kojo Acquah and the mass spectrometry facility within the Department of Chemistry at Western Michigan University. This work was supported by the NIH grant R15GM163283, and startup funds and a grant from the Faculty Research and Creative Activities Award, Western Michigan University.

---

*Supplementary Figure 1*

*S. flexneri* expression of PDEs in their respective mutant strains does not significantly alter growth. (A) OD_600_ of *S. flexneri* strains was measured over time. (B) The growth rate of each strain during exponential growth was calculated.

*Supplementary Figure 2*

*S. flexneri* Δ*pde* mutant strains expressing their respective genes showed a significant reduction in biofilm formation. # indicates which strains were statically significant to WT.

*Supplementary Figure 3*

Henle-407 cells infected with *Shigella* PDE complementation strains did not show any statistical significance compared to WT when the invasion assay was performed. # indicates which strains were statically significant to WT.

*Supplementary Figure 4*

Δ*pde*B + *pde*B, Δ*pde*D + *pde*D, Δ*pde*G + *pde*G and Δ*pde*K + *pde*K all showed decreases in plaque size. Δ*pde*H + *pde*H and Δ*pde*N + *pde*N had larger plaques sizes compared to WT. # indicates which strains were statically significant to WT.

*Supplementary Figure 5*

Quantification of c-di-GMP levels in *Shigella*’s c-di-GMP specific PDEs mutant strains using microscopy. Graphs A and B represents biological replicates 2 and 3 replicates.

*Supplementary Figure 6*

Δ*pde*B pathogenicity genes upregulated when compared to WT.

*Supplementary Figure 7*

Pictures of *S. flexneri* plates (A) WT, (B) CFS100 and (C) *ΔfadD* showing the differences colony color.

*Supplementary Figure 8*

Showing differences in FAs in *S.flexneri* WT, Δ*pdeB,* Δ*pde+B* and *ΔfadD* when grown to stationary phase. Statistical significance between WT and *ΔfadD* was seen for (ii) C15 and (iv) C16 FAs.

## References

1. Aslam A, Hashmi M, Okafor C. Shigellosis [Internet]. StatPearls Publishing; 2024. Available from: https://www.ncbi.nlm.nih.gov/books/NBK482337/

2. Mason LCE, Charles H, Thorley K, Chong CE, De Silva PM, Jenkins C, et al. The re-emergence of sexually transmissible multidrug resistant Shigella flexneri 3a, England, United Kingdom. npj Antimicrob Resist. 2024 Aug 2;2(1):20. doi:10.1038/s44259-024-00038-3

3. Muzembo BA, Kitahara K, Mitra D, Ohno A, Khatiwada J, Dutta S, et al. Burden of *Shigella* in South Asia: a systematic review and meta-analysis. Journal of Travel Medicine. 2023 Feb 18;30(1):taac132. doi:10.1093/jtm/taac132

4. Kotloff KL, Riddle MS, Platts-Mills JA, Pavlinac P, Zaidi AKM. Shigellosis. The Lancet. 2018 Feb;391(10122):801–12. doi:10.1016/S0140-6736(17)33296-8

5. Magalis BR, Salemi M. Molecular epidemiology of foodborne pathogens. In: Foodborne Infections and Intoxications [Internet]. Elsevier; 2021 [cited 2025 Aug 28]. p. 47–62. Available from: https://linkinghub.elsevier.com/retrieve/pii/B9780128195192000074 doi:10.1016/B978-0-12-819519-2.00007-4

6. Baker S, The HC. Recent insights into Shigella: a major contributor to the global diarrhoeal disease burden. Current Opinion in Infectious Diseases. 2018 Oct;31(5):449–54. doi:10.1097/QCO.0000000000000475

7. Zaidi MB, Estrada-García T. Shigella: A Highly Virulent and Elusive Pathogen. Curr Trop Med Rep. 2014 Mar 25. doi:10.1007/s40475-014-0019-6

8. Giersing BK, Isbrucker R, Kaslow DC, Cavaleri M, Baylor N, Maiga D, et al. Clinical and regulatory development strategies for Shigella vaccines intended for children younger than 5 years in low-income and middle-income countries. The Lancet Global Health. 2023 Nov;11(11):e1819–26. doi:10.1016/S2214-109X(23)00421-7

9. Desalegn G, Abrahamson C, Ross Turbyfill K, Pill-Pepe L, Bautista L, Tamilselvi CS, et al. A broad spectrum Shigella vaccine based on VirG53–353 multiepitope region produced in a cell-free system. npj Vaccines. 2025 Jan 13;10(1):6. doi:10.1038/s41541-025-01064-6

10. Undale VR, Gupta S, Lakhadive K. Novel Targets for Antimicrobials. tjps. 2020 Oct 1;17(5):565–75. doi:10.4274/tjps.galenos.2020.90197

11. Cardona ST, Rahman ASMZ, Novomisky Nechcoff J. Innovative perspectives on the discovery of small molecule antibiotics. npj Antimicrob Resist. 2025 Mar 13;3(1):19. doi:10.1038/s44259-025-00089-0

12. Schroeder GN, Hilbi H. Molecular Pathogenesis of *Shigella* spp.: Controlling Host Cell Signaling, Invasion, and Death by Type III Secretion. Clin Microbiol Rev. 2008 Jan;21(1):134–56. doi:10.1128/CMR.00032-07

13. Chattaway MA, Schaefer U, Tewolde R, Dallman TJ, Jenkins C. Identification of Escherichia coli and Shigella Species from Whole-Genome Sequences. Ledeboer NA, editor. J Clin Microbiol. 2017 Feb;55(2):616–23. doi:10.1128/JCM.01790-16

14. Devanga Ragupathi NK, Muthuirulandi Sethuvel DP, Inbanathan FY, Veeraraghavan B. Accurate differentiation of Escherichia coli and Shigella serogroups: challenges and strategies. New Microbes and New Infections. 2018 Jan;21:58–62. doi:10.1016/j.nmni.2017.09.003

15. Day WA, Fernández RE, Maurelli AT. Pathoadaptive Mutations That Enhance Virulence: Genetic Organization of the *cadA* Regions of *Shigella* spp. Barbieri JT, editor. Infect Immun. 2001 Dec;69(12):7471–80. doi:10.1128/IAI.69.12.7471-7480.2001

16. Niu C, Yang J, Liu H, Cui Y, Xu H, Wang R, et al. Role of the virulence plasmid in acid resistance of Shigella flexneri. Sci Rep. 2017 Apr 25;7(1):46465. doi:10.1038/srep46465

17. Lan R, Reeves PR. Escherichia coli in disguise: molecular origins of Shigella. Microbes and Infection. 2002 Sep;4(11):1125–32. doi:10.1016/S1286-4579(02)01637-4

18. Hershberg R, Tang H, Petrov DA. Reduced selection leads to accelerated gene loss in Shigella. Genome Biol. 2007 Aug 8;8(8):R164. doi:10.1186/gb-2007-8-8-r164

19. Gorden J, Small PL. Acid resistance in enteric bacteria. Infect Immun. 1993 Jan;61(1):364–7. doi:10.1128/iai.61.1.364-367.1993

20. Martinić M, Hoare A, Contreras I, Álvarez SA. Contribution of the Lipopolysaccharide to Resistance of Shigella flexneri 2a to Extreme Acidity. Adler B, editor. PLoS ONE. 2011 Oct 3;6(10):e25557. doi:10.1371/journal.pone.0025557

21. Schnupf P, Sansonetti PJ. *Shigella* Pathogenesis: New Insights through Advanced Methodologies. Cossart P, Roy CR, Sansonetti P, editors. Microbiol Spectr. 2019 Apr 12;7(2):7.2.28. doi:10.1128/microbiolspec.BAI-0023-2019

22. Chiang IL, Wang Y, Fujii S, Muegge BD, Lu Q, Tarr PI, et al. Biofilm Formation and Virulence of Shigella flexneri Are Modulated by pH of Gastrointestinal Tract. Brodsky IE, editor. Infect Immun. 2021 Oct 15;89(11):e00387–21. doi:10.1128/IAI.00387-21

23. Philpott DJ, Edgeworth JD, Sansonetti PJ. The pathogenesis of *Shigella flexneri* infection: lessons from *in vitro* and in vivo studies. Smith H, Dorman CJ, Dougan G, Holden DW, Dougan G, Williams P, editors. Phil Trans R Soc Lond B. 2000 May 28;355(1397):575–86. doi:10.1098/rstb.2000.0599

24. Carayol N, Tran Van Nhieu G. The Inside Story of Shigella Invasion of Intestinal Epithelial Cells. Cold Spring Harbor Perspectives in Medicine. 2013 Oct 1;3(10):a016717–a016717. doi:10.1101/cshperspect.a016717

25. Nasser A, Mosadegh M, Azimi T, Shariati A. Molecular mechanisms of Shigella effector proteins: a common pathogen among diarrheic pediatric population. Mol Cell Pediatr. 2022 Jun 19;9(1):12. doi:10.1186/s40348-022-00145-z

26. Agaisse H. Molecular and Cellular Mechanisms of Shigella flexneri Dissemination. Front Cell Infect Microbiol. 2016 Mar 11;6. doi:10.3389/fcimb.2016.00029

27. Carayol N, Tran Van Nhieu G. Tips and tricks about Shigella invasion of epithelial cells. Current Opinion in Microbiology. 2013 Feb;16(1):32–7. doi:10.1016/j.mib.2012.11.010

28. Almblad H, Randall TE, Liu F, Leblanc K, Groves RA, Kittichotirat W, et al. Bacterial cyclic diguanylate signaling networks sense temperature. Nat Commun. 2021 Mar 31;12(1):1986. doi:10.1038/s41467-021-22176-2

29. Randall TE, Eckartt K, Kakumanu S, Price-Whelan A, Dietrich LEP, Harrison JJ. Sensory Perception in Bacterial Cyclic Diguanylate Signal Transduction. O’Toole G, editor. J Bacteriol. 2022 Feb 15;204(2):e00433–21. doi:10.1128/jb.00433-21

30. Jenal U, Reinders A, Lori C. Cyclic di-GMP: second messenger extraordinaire. Nat Rev Microbiol. 2017 May;15(5):271–84. doi:10.1038/nrmicro.2016.190

31. Martín-Rodríguez AJ, Higdon SM, Thorell K, Tellgren-Roth C, Sjöling Å, Galperin MY, et al. Comparative Genomics of Cyclic di-GMP Metabolism and Chemosensory Pathways in Shewanella algae Strains: Novel Bacterial Sensory Domains and Functional Insights into Lifestyle Regulation. Kovács ÁT, editor. mSystems. 2022 Apr 26;7(2):e01518–21. doi:10.1128/msystems.01518-21

32. Wang Z, Song L, Liu X, Shen X, Li X. Bacterial second messenger c-di-GMP: Emerging functions in stress resistance. Microbiological Research. 2023 Mar;268:127302. doi:10.1016/j.micres.2023.127302

33. Zhan X, Zhang K, Wang C, Fan Q, Tang X, Zhang X, et al. A c-di-GMP signaling module controls responses to iron in Pseudomonas aeruginosa. Nat Commun. 2024 Feb 29;15(1):1860. doi:10.1038/s41467-024-46149-3

34. Hengge R. Principles of c-di-GMP signalling in bacteria. Nat Rev Microbiol. 2009 Apr;7(4):263–73. doi:10.1038/nrmicro2109

35. Ojha R, Krug S, Jones P, Koestler BJ. Intact and mutated Shigella diguanylate cyclases increase c-di-GMP. Journal of Biological Chemistry. 2024 Aug;300(8):107525. doi:10.1016/j.jbc.2024.107525

36. Ojha R, Dittmar AA, Severin GB, Koestler BJ. Shigella flexneri Diguanylate Cyclases Regulate Virulence. Galperin MY, editor. J Bacteriol. 2021 Nov 5;203(23):e00242–21. doi:10.1128/JB.00242-21

37. Herbst S, Lorkowski M, Sarenko O, Nguyen TKL, Jaenicke T, Hengge R. Transmembrane redox control and proteolysis of PdeC, a novel type of c-di- GMP phosphodiesterase. The EMBO Journal. 2018 Apr 13;37(8):e97825. doi:10.15252/embj.201797825

38. Hengge R, Galperin MY, Ghigo JM, Gomelsky M, Green J, Hughes KT, et al. Systematic Nomenclature for GGDEF and EAL Domain-Containing Cyclic Di-GMP Turnover Proteins of Escherichia coli. O’Toole GA, editor. J Bacteriol. 2016 Jan;198(1):7–11. doi:10.1128/JB.00424-15

39. Hengge R, Gründling A, Jenal U, Ryan R, Yildiz F. Bacterial Signal Transduction by Cyclic Di-GMP and Other Nucleotide Second Messengers. O’Toole GA, editor. J Bacteriol. 2016 Jan;198(1):15–26. doi:10.1128/JB.00331-15

40. Schmidt AJ, Ryjenkov DA, Gomelsky M. The Ubiquitous Protein Domain EAL Is a Cyclic Diguanylate-Specific Phosphodiesterase: Enzymatically Active and Inactive EAL Domains. J Bacteriol. 2005 Jul;187(14):4774–81. doi:10.1128/JB.187.14.4774-4781.2005

41. Chatterjee D, Cooley RB, Boyd CD, Mehl RA, O’Toole GA, Sondermann H. Mechanistic insight into the conserved allosteric regulation of periplasmic proteolysis by the signaling molecule cyclic-di-GMP. eLife. 2014 Sep 2;3:e03650. doi:10.7554/eLife.03650

42. Richter AM, Possling A, Malysheva N, Yousef KP, Herbst S, Von Kleist M, et al. Local c-di-GMP Signaling in the Control of Synthesis of the E. coli Biofilm Exopolysaccharide pEtN-Cellulose. Journal of Molecular Biology. 2020 Jul;432(16):4576–95. doi:10.1016/j.jmb.2020.06.006

43. Simm R, Morr M, Kader A, Nimtz M, Römling U. GGDEF and EAL domains inversely regulate cyclic di-GMP levels and transition from sessility to motility. Molecular Microbiology. 2004 Aug;53(4):1123–34. doi:10.1111/j.1365-2958.2004.04206.x

44. Reinders A, Hee CS, Ozaki S, Mazur A, Boehm A, Schirmer T, et al. Expression and Genetic Activation of Cyclic Di-GMP-Specific Phosphodiesterases in Escherichia coli. Zhulin IB, editor. J Bacteriol. 2016 Feb;198(3):448–62. doi:10.1128/JB.00604-15

45. The UniProt Consortium, Bateman A, Martin MJ, Orchard S, Magrane M, Adesina A, et al. UniProt: the Universal Protein Knowledgebase in 2025. Nucleic Acids Research. 2025 Jan 6;53(D1):D609–17. doi:10.1093/nar/gkae1010

46. Datsenko KA, Wanner BL. One-step inactivation of chromosomal genes in *Escherichia coli* K-12 using PCR products. Proc Natl Acad Sci USA. 2000 Jun 6;97(12):6640–5. doi:10.1073/pnas.120163297

47. Liu C, Shi R, Jensen MS, Zhu J, Liu J, Liu X, et al. The global regulation of c-di-GMP and cAMP in bacteria. mLife. 2024 Mar;3(1):42–56. doi:10.1002/mlf2.12104

48. Kim HK, Harshey RM. A Diguanylate Cyclase Acts as a Cell Division Inhibitor in a Two-Step Response to Reductive and Envelope Stresses. Gottesman S, editor. mBio. 2016 Sep 7;7(4):e00822–16. doi:10.1128/mBio.00822-16

49. Wei J, Goldberg MB, Burland V, Venkatesan MM, Deng W, Fournier G, et al. Complete Genome Sequence and Comparative Genomics of *Shigella flexneri* Serotype 2a Strain 2457T. Infect Immun. 2003 May;71(5):2775–86. doi:10.1128/IAI.71.5.2775-2786.2003

50. Nickerson KP, Chanin RB, Sistrunk JR, Rasko DA, Fink PJ, Barry EM, et al. Analysis of Shigella flexneri Resistance, Biofilm Formation, and Transcriptional Profile in Response to Bile Salts. McCormick B, editor. Infect Immun. 2017 Jun;85(6):e01067–16. doi:10.1128/IAI.01067-16

51. Zouhir S, Abidi W, Caleechurn M, Krasteva PV. Structure and Multitasking of the c-di-GMP-Sensing Cellulose Secretion Regulator BcsE. Harwood CS, editor. mBio. 2020 Aug 25;11(4):e01303–20. doi:10.1128/mBio.01303-20

52. Sen T, Verma NK. YfiB: An Outer Membrane Protein Involved in the Virulence of Shigella flexneri. Microorganisms. 2022 Mar 18;10(3):653. doi:10.3390/microorganisms10030653

53. Koestler BJ, Ward CM, Payne SM. *Shigella* Pathogenesis Modeling with Tissue Culture Assays. CP Microbiology. 2018 Aug;50(1):e57. doi:10.1002/cpmc.57

54. Ross ND, Chin AL, Pannuri A, Doore SM. Bacterial lysis or survival after infection with phage Sf14 depends on combined nutrient and temperature conditions. Onohuean H, editor. PLoS ONE. 2025 Mar 25;20(3):e0319836. doi:10.1371/journal.pone.0319836

55. Koestler BJ, Ward CM, Fisher CR, Rajan A, Maresso AW, Payne SM. Human Intestinal Enteroids as a Model System of *Shigella* Pathogenesis. Young VB, editor. Infect Immun. 2019 Apr;87(4):e00733–18. doi:10.1128/IAI.00733-18

56. Brotcke Zumsteg A, Goosmann C, Brinkmann V, Morona R, Zychlinsky A. IcsA Is a Shigella flexneri Adhesin Regulated by the Type III Secretion System and Required for Pathogenesis. Cell Host & Microbe. 2014 Apr;15(4):435–45. doi:10.1016/j.chom.2014.03.001

57. Mauricio RPM, Jeffries CM, Svergun DI, Deane JE. The Shigella Virulence Factor IcsA Relieves N-WASP Autoinhibition by Displacing the Verprolin Homology/Cofilin/Acidic (VCA) Domain. Journal of Biological Chemistry. 2017 Jan;292(1):134–45. doi:10.1074/jbc.M116.758003

58. Anderson WA, Dippel AB, Maiden MM, Waters CM, Hammond MC. Chemiluminescent sensors for quantitation of the bacterial second messenger cyclic di-GMP. In: Methods in Enzymology [Internet]. Elsevier; 2020 [cited 2025 Aug 28]. p. 83–104. Available from: https://linkinghub.elsevier.com/retrieve/pii/S0076687920301324 doi:10.1016/bs.mie.2020.04.004

59. Koestler BJ, Waters CM. Exploring Environmental Control of Cyclic di-GMP Signaling in Vibrio cholerae by Using the *Ex Vivo* Lysate Cyclic di-GMP Assay (TELCA). Appl Environ Microbiol. 2013 Sep;79(17):5233–41. doi:10.1128/AEM.01596-13

60. Nieto V, Partridge JD, Severin GB, Lai RZ, Waters CM, Parkinson JS, et al. Under Elevated c-di-GMP in Escherichia coli, YcgR Alters Flagellar Motor Bias and Speed Sequentially, with Additional Negative Control of the Flagellar Regulon via the Adaptor Protein RssB. O’Toole G, editor. J Bacteriol. 2019 Dec 6;202(1). doi:10.1128/JB.00578-19

61. Sarenko O, Klauck G, Wilke FM, Pfiffer V, Richter AM, Herbst S, et al. More than Enzymes That Make or Break Cyclic Di-GMP—Local Signaling in the Interactome of GGDEF/EAL Domain Proteins of *Escherichia coli*. Shuman HA, editor. mBio. 2017 Nov 8;8(5):e01639–17. doi:10.1128/mBio.01639-17

62. Zhou H, Zheng C, Su J, Chen B, Fu Y, Xie Y, et al. Characterization of a natural triple-tandem c-di-GMP riboswitch and application of the riboswitch-based dual-fluorescence reporter. Sci Rep. 2016 Feb 19;6(1):20871. doi:10.1038/srep20871

63. UK Standards for Microbiology Investigations Identification of Shigella species. [Internet]. Standards Unit, UK Health Security Agency; 2025. Available from: https://www.rcpath.org/static/e2671e77-205a-4ef2-91bd00f25f4192fe/uk-smi-id-20i4-1-identification-of-shigella-species-july-2025-pdf.pdf

64. Sun QX, Huang M, Zhang JY, Zeng X, Zhang CC. Control of Cell Size by c-di-GMP Requires a Two-Component Signaling System in the Cyanobacterium *Anabaena* sp. Strain PCC 7120. Wang L, editor. Microbiol Spectr. 2023 Feb 14;11(1):e04228-22. doi:10.1128/spectrum.04228-22

65. Love MI, Huber W, Anders S. Moderated estimation of fold change and dispersion for RNA-seq data with DESeq2. Genome Biol. 2014 Dec 5;15(12):550. doi:10.1186/s13059-014-0550-8

66. Niebuhr K, Jouihri N, Allaoui A, Gounon P, Sansonetti PJ, Parsot C. IpgD, a protein secreted by the type III secretion machinery of *Shigella flexneri*, is chaperoned by IpgE and implicated in entry focus formation. Molecular Microbiology. 2000 Oct;38(1):8–19. doi:10.1046/j.1365-2958.2000.02041.x

67. Tran Van Nhieu G, Latour-Lambert P, Enninga J. Modification of phosphoinositides by the Shigella effector IpgD during host cell infection. Front Cell Infect Microbiol. 2022 Oct 27;12:1012533. doi:10.3389/fcimb.2022.1012533

68. Janßen H, Steinbüchel A. Fatty acid synthesis in Escherichia coli and its applications towards the production of fatty acid based biofuels. Biotechnol Biofuels. 2014;7(1):7. doi:10.1186/1754-6834-7-7

69. Janßen H, Steinbüchel A. Fatty acid synthesis in Escherichia coli and its applications towards the production of fatty acid based biofuels. Biotechnol Biofuels. 2014;7(1):7. doi:10.1186/1754-6834-7-7

70. Trirocco R, Pasqua M, Tramonti A, Grossi M, Colonna B, Paiardini A, et al. Fatty Acids Abolish *Shigella* Virulence by Inhibiting Its Master Regulator, VirF. Cascales E, editor. Microbiol Spectr. 2023 Jun 15;11(3):e00778–23. doi:10.1128/spectrum.00778-23

71. Politz M, Lennen R, Pfleger B. Quantification of Bacterial Fatty Acids by Extraction and Methylation. BIO-PROTOCOL. 2013;3(21). doi:10.21769/BioProtoc.950

72. Vejzovic D, Schwaiger T, Topciu A, Petrowitsch L, Arnautovic A, Malanovic N. Bacterial cell fate under stress: lipid remodeling and antimicrobial peptide attack. npj Antimicrob Resist. 2026 Apr 2;4(1):22. doi:10.1038/s44259-026-00195-7

73. MacDermott-Opeskin HI, Gupta V, O’Mara ML. Lipid-mediated antimicrobial resistance: a phantom menace or a new hope? Biophys Rev. 2022 Feb;14(1):145–62. doi:10.1007/s12551-021-00912-8

74. Römling U, Galperin MY, Gomelsky M. Cyclic di-GMP: the First 25 Years of a Universal Bacterial Second Messenger. Microbiol Mol Biol Rev. 2013 Mar;77(1):1–52. doi:10.1128/MMBR.00043-12

75. Boyd CD, O’Toole GA. Second Messenger Regulation of Biofilm Formation: Breakthroughs in Understanding c-di-GMP Effector Systems. Annual Review of Cell and Developmental Biology. 2012 Nov 10;28(1):439–62. doi:10.1146/annurev-cellbio-101011-155705

76. Park S, Sauer K. Controlling Biofilm Development Through Cyclic di-GMP Signaling. In: Filloux A, Ramos JL, editors. Pseudomonas aeruginosa [Internet]. Cham: Springer International Publishing; 2022 [cited 2025 Sep 13]. p. 69–94. (Advances in Experimental Medicine and Biology). Available from: https://link.springer.com/10.1007/978-3-031-08491-1_3 doi:10.1007/978-3-031-08491-1_3

77. Hwang Y, Perez M, Holzel R, Harshey RM. c-di-GMP is required for swarming in *E. coli*, producing colanic acid that acts as surfactant. Harwood CS, editor. mBio. 2025 Jun 11;16(6):e00916–25. doi:10.1128/mbio.00916-25

78. Sakellaris H, Hannink NK, Rajakumar K, Bulach D, Hunt M, Sasakawa C, et al. Curli Loci of *Shigella* spp. O’Brien AD, editor. Infect Immun. 2000 Jun;68(6):3780–3. doi:10.1128/IAI.68.6.3780-3783.2000

79. Römling U, Galperin MY. Bacterial cellulose biosynthesis: diversity of operons, subunits, products, and functions. Trends in Microbiology. 2015 Sep;23(9):545–57. doi:10.1016/j.tim.2015.05.005

80. Chanin RB, Nickerson KP, Llanos-Chea A, Sistrunk JR, Rasko DA, Kumar DKV, et al. *Shigella flexneri* Adherence Factor Expression in *In Vivo* -Like Conditions. Ellermeier CD, editor. mSphere. 2019 Dec 18;4(6):e00751–19. doi:10.1128/mSphere.00751-19

81. Serra DO, Hengge R. A c-di-GMP-Based Switch Controls Local Heterogeneity of Extracellular Matrix Synthesis which Is Crucial for Integrity and Morphogenesis of Escherichia coli Macrocolony Biofilms. Journal of Molecular Biology. 2019 Nov;431(23):4775–93. doi:10.1016/j.jmb.2019.04.001

82. Valentini M, Filloux A. Multiple Roles of c-di-GMP Signaling in Bacterial Pathogenesis. Annu Rev Microbiol. 2019 Sep 8;73(1):387–406. doi:10.1146/annurev-micro-020518-115555

83. Conner JG, Zamorano-Sánchez D, Park JH, Sondermann H, Yildiz FH. The ins and outs of cyclic di-GMP signaling in Vibrio cholerae. Current Opinion in Microbiology. 2017 Apr;36:20–9. doi:10.1016/j.mib.2017.01.002

84. Rosen DA, Twentyman J, Hunstad DA. High Levels of Cyclic Di-GMP in Klebsiella pneumoniae Attenuate Virulence in the Lung. Payne SM, editor. Infect Immun. 2018 Feb;86(2):e00647–17. doi:10.1128/IAI.00647-17

85. Li S, Sun H, Li J, Zhao Y, Wang R, Xu L, et al. Autoinducer-2 and bile salts induce c-di-GMP synthesis to repress the T3SS via a T3SS chaperone. Nat Commun. 2022 Nov 5;13(1):6684. doi:10.1038/s41467-022-34607-9

86. Lamprokostopoulou A, Monteiro C, Rhen M, Römling U. Cyclic di-GMP signalling controls virulence properties of *Salmonella enterica* serovar Typhimurium at the mucosal lining. Environmental Microbiology. 2010 Jan;12(1):40–53. doi:10.1111/j.1462-2920.2009.02032.x

87. Ling X, Liu X, Wang K, Guo M, Ou Y, Li D, et al. Lsr2 acts as a cyclic di-GMP receptor that promotes keto-mycolic acid synthesis and biofilm formation in mycobacteria. Nat Commun. 2024 Jan 24;15(1):695. doi:10.1038/s41467-024-44774-6

88. Lefebvre C, Boulon R, Ducoux M, Gavalda S, Laval F, Jamet S, et al. HadD, a novel fatty acid synthase type II protein, is essential for alpha- and epoxy-mycolic acid biosynthesis and mycobacterial fitness. Sci Rep. 2018 Apr 16;8(1):6034. doi:10.1038/s41598-018-24380-5

89. Trirocco R, Pasqua M, Tramonti A, Grossi M, Colonna B, Paiardini A, et al. Fatty Acids Abolish *Shigella* Virulence by Inhibiting Its Master Regulator, VirF. Cascales E, editor. Microbiol Spectr. 2023 Jun 15;11(3):e00778–23. doi:10.1128/spectrum.00778-23

90. Giangrossi M, Giuliodori AM, Tran CN, Amici A, Marchini C, Falconi M. VirF Relieves the Transcriptional Attenuation of the Virulence Gene icsA of Shigella flexneri Affecting the icsA mRNA–RnaG Complex Formation. Front Microbiol. 2017 Apr 18;8. doi:10.3389/fmicb.2017.00650

91. Gerson TM, Ott AM, Karney MMA, Socea JN, Ginete DR, Iyer LM, et al. VirB, a key transcriptional regulator of *Shigella* virulence, requires a CTP ligand for its regulatory activities. Babitzke P, editor. mBio. 2023 Oct 31;14(5):e01519–23. doi:10.1128/mbio.01519-23

92. Di Martino ML, Falconi M, Micheli G, Colonna B, Prosseda G. The Multifaceted Activity of the VirF Regulatory Protein in the Shigella Lifestyle. Front Mol Biosci. 2016 Sep 29;3. doi:10.3389/fmolb.2016.00061

93. Murakami R, Xiao J zhong, Hara KY, Odamaki T, Kikukawa H. Comprehensive analysis of cell-associated fatty acids in *Bifidobacterium* strains. Bioscience, Biotechnology, and Biochemistry. 2025 Nov 25;89(12):1700–5. doi:10.1093/bbb/zbaf144

94. Lucas RL, Lostroh CP, DiRusso CC, Spector MP, Wanner BL, Lee CA. Multiple Factors Independently Regulate *hilA* and Invasion Gene Expression in *Salmonella enterica* Serovar Typhimurium. J Bacteriol. 2000 Apr;182(7):1872–82. doi:10.1128/JB.182.7.1872-1882.2000

95. Ray S, Chatterjee E, Chatterjee A, Paul K, Chowdhury R. A *fadD* mutant of *Vibrio cholerae* Is Impaired in the Production of Virulence Factors and Membrane Localization of the Virulence Regulatory Protein TcpP. Payne SM, editor. Infect Immun. 2011 Jan;79(1):258–66. doi:10.1128/IAI.00663-10

96. Westfall CS, Levin PA. Bacterial Cell Size: Multifactorial and Multifaceted. Annu Rev Microbiol. 2017 Sep 8;71(1):499–517. doi:10.1146/annurev-micro-090816-093803

97. Kaczmarczyk A, Hempel AM, Von Arx C, Böhm R, Dubey BN, Nesper J, et al. Precise timing of transcription by c-di-GMP coordinates cell cycle and morphogenesis in Caulobacter. Nat Commun. 2020 Feb 10;11(1):816. doi:10.1038/s41467-020-14585-6

98. Malone JG, Jaeger T, Spangler C, Ritz D, Spang A, Arrieumerlou C, et al. YfiBNR Mediates Cyclic di-GMP Dependent Small Colony Variant Formation and Persistence in Pseudomonas aeruginosa. Roy CR, editor. PLoS Pathog. 2010 Mar 12;6(3):e1000804. doi:10.1371/journal.ppat.1000804

99. Shin S, Yu J, Tae H, Zhao Y, Jiang D, Qiao Y, et al. Exploring the Membrane-Active Interactions of Antimicrobial Long-Chain Fatty Acids Using a Supported Lipid Bilayer Model for Gram-Positive Bacterial Membranes. ACS Appl Mater Interfaces. 2024 Oct 23;16(42):56705–17. doi:10.1021/acsami.4c11158

100. Nicastro GG, Kaihami GH, Pereira TO, Meireles DA, Groleau M, Déziel E, et al. C yclic-di-GMP levels affect *P seudomonas aeruginosa* fitness in the presence of imipenem. Environmental Microbiology. 2014 May;16(5):1321–33. doi:10.1111/1462-2920.12422

101. Gupta K, Liao J, Petrova OE, Cherny KE, Sauer K. Elevated levels of the second messenger c-di- GMP contribute to antimicrobial resistance of *P seudomonas aeruginosa*. Molecular Microbiology. 2014 May;92(3):488–506. doi:10.1111/mmi.12587

102. Sidik S, Kottwitz H, Benjamin J, Ryu J, Jarrar A, Garduno R, et al. A *Shigella flexneri* Virulence Plasmid Encoded Factor Controls Production of Outer Membrane Vesicles. G3 Genes|Genomes|Genetics. 2014 Dec 1;4(12):2493–503. doi:10.1534/g3.114.014381

103. Baba T, Ara T, Hasegawa M, Takai Y, Okumura Y, Baba M, et al. Construction of *Escherichia coli* K-12 in-frame, single-gene knockout mutants: the Keio collection. Molecular Systems Biology. 2006 Jan;2(1):2006.0008. doi:10.1038/msb4100050

104. Saragliadis A, Trunk T, Leo JC. Producing Gene Deletions in Escherichia coli by P1 Transduction with Excisable Antibiotic Resistance Cassettes. J Vis Exp. 2018 Sep 1;(139):58267. doi:10.3791/58267 PubMed PMID: 30222159; PubMed Central PMCID: PMC6235078.

105. Bushnell B. BBMap: A Fast, Accurate, Splice-Aware Aligner. In. 2014.

106. Li H. Aligning sequence reads, clone sequences and assembly contigs with BWA-MEM [Internet]. arXiv; 2013 [cited 2026 Apr 11]. Available from: https://arxiv.org/abs/1303.3997doi:10.48550/ARXIV.1303.3997

107. Li H, Durbin R. Fast and accurate short read alignment with Burrows–Wheeler transform. Bioinformatics. 2009 Jul 15;25(14):1754–60. doi:10.1093/bioinformatics/btp324

108. Li H, Handsaker B, Wysoker A, Fennell T, Ruan J, Homer N, et al. The Sequence Alignment/Map format and SAMtools. Bioinformatics. 2009 Aug 15;25(16):2078–9. doi:10.1093/bioinformatics/btp352

109. Liao Y, Smyth GK, Shi W. featureCounts: an efficient general purpose program for assigning sequence reads to genomic features. Bioinformatics. 2014 Apr 1;30(7):923–30. doi:10.1093/bioinformatics/btt656

110. Gentleman RC, Carey VJ, Bates DM, Bolstad B, Dettling M, Dudoit S, et al. Bioconductor: open software development for computational biology and bioinformatics. Genome Biol. 2004 Sep 15;5(10):R80. doi:10.1186/gb-2004-5-10-r80

111. Blighe K, Rana S, Lewis M. EnhancedVolcano: publication-ready volcano plots with enhanced colouring and labeling [Internet]. 2026 Apr 28. Available from: https://bioconductor.org/packages/release/bioc/vignettes/EnhancedVolcano/inst/doc/EnhancedVolcano.html

112. Yu G, Wang LG, Han Y, He QY. clusterProfiler: an R Package for Comparing Biological Themes Among Gene Clusters. OMICS: A Journal of Integrative Biology. 2012 May 1;16(5):284–7. doi:10.1089/omi.2011.0118

113. Kanehisa M, Furumichi M, Tanabe M, Sato Y, Morishima K. KEGG: new perspectives on genomes, pathways, diseases and drugs. Nucleic Acids Res. 2017 Jan 4;45(D1):D353–61. doi:10.1093/nar/gkw1092

114. Ren J, Mozurkewich E, Sen A, Vahratian A, Ferreri T, Morse A, et al. Total Serum Fatty Acid Analysis by GC-MS: Assay Validation and Serum Sample Stability. CPA. 2013 Jul 1;9(4):331–9. doi:10.2174/1573412911309040002

115. Hudzicki J. Kirby-Bauer Disk Diffusion Susceptibility Test Protocol [Internet]. American Society for Microbiology; 2009. Available from: https://asm.org/getattachment/2594ce26-bd44-47f6-8287-0657aa9185ad/kirby-bauer-disk-diffusion-

