## Supplementary figures and images for "*Shigella*’s c-di-GMP specific PDEs Modulate Biofilm and Virulence Phenotypes"

### Fig S1

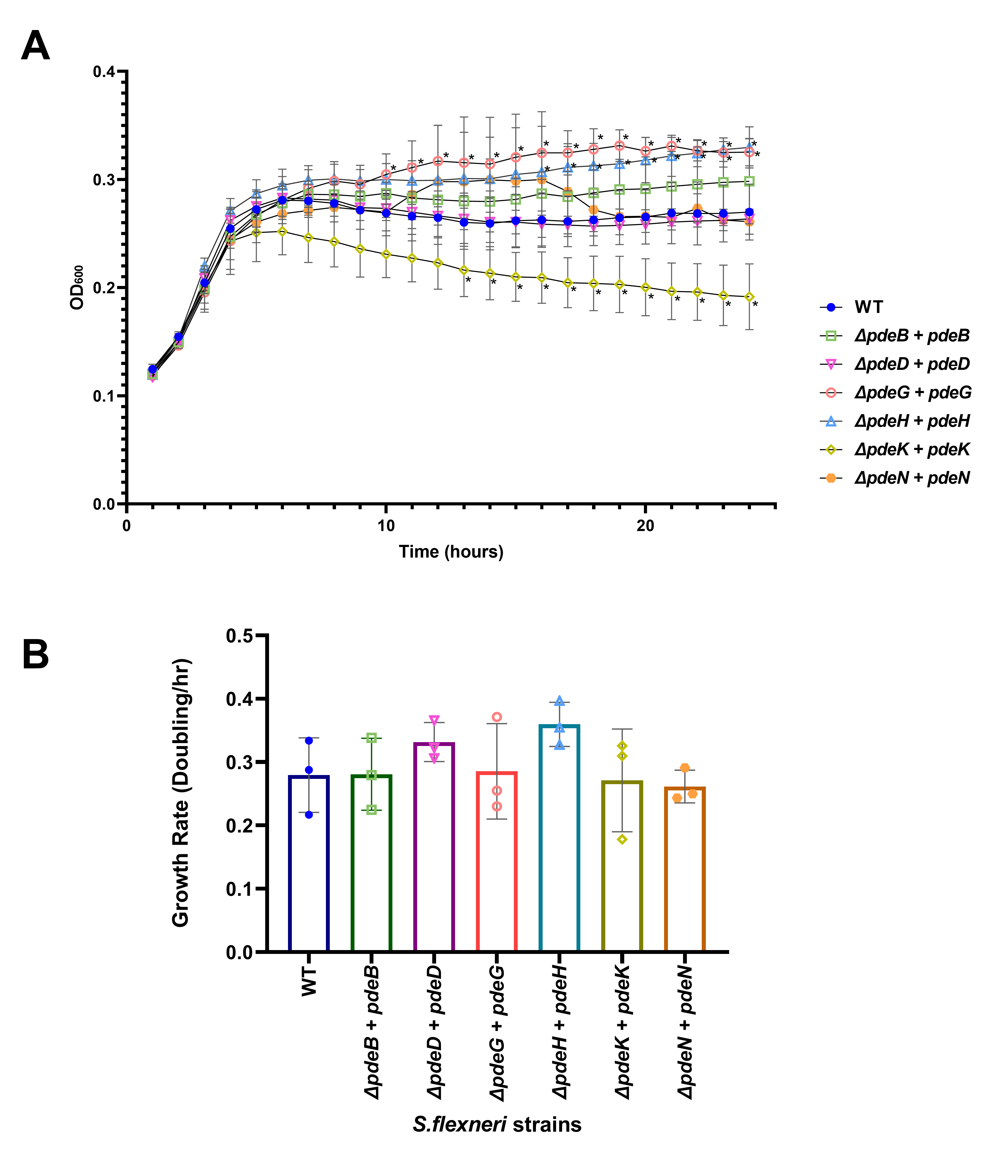

### Fig S2

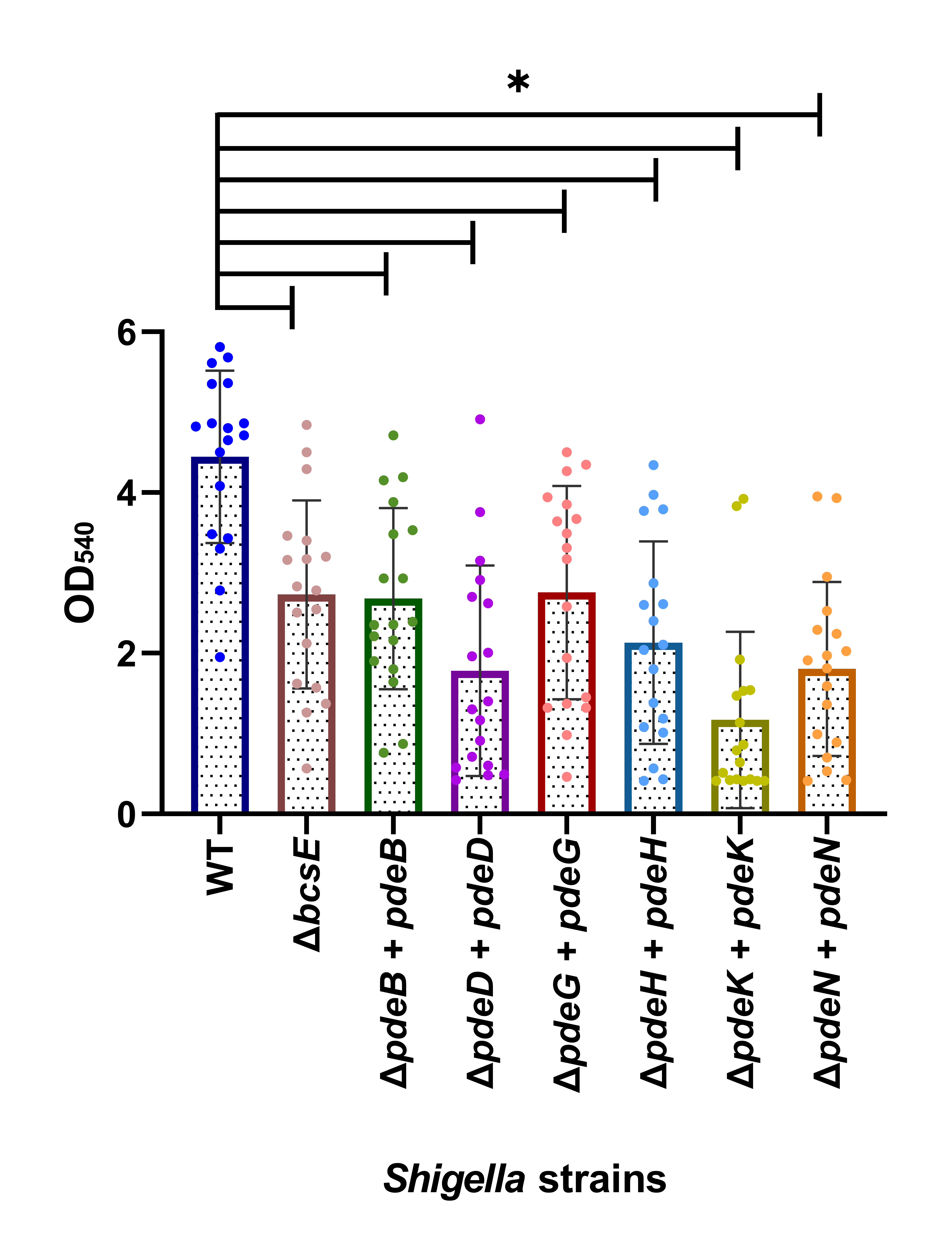

### Fig S3

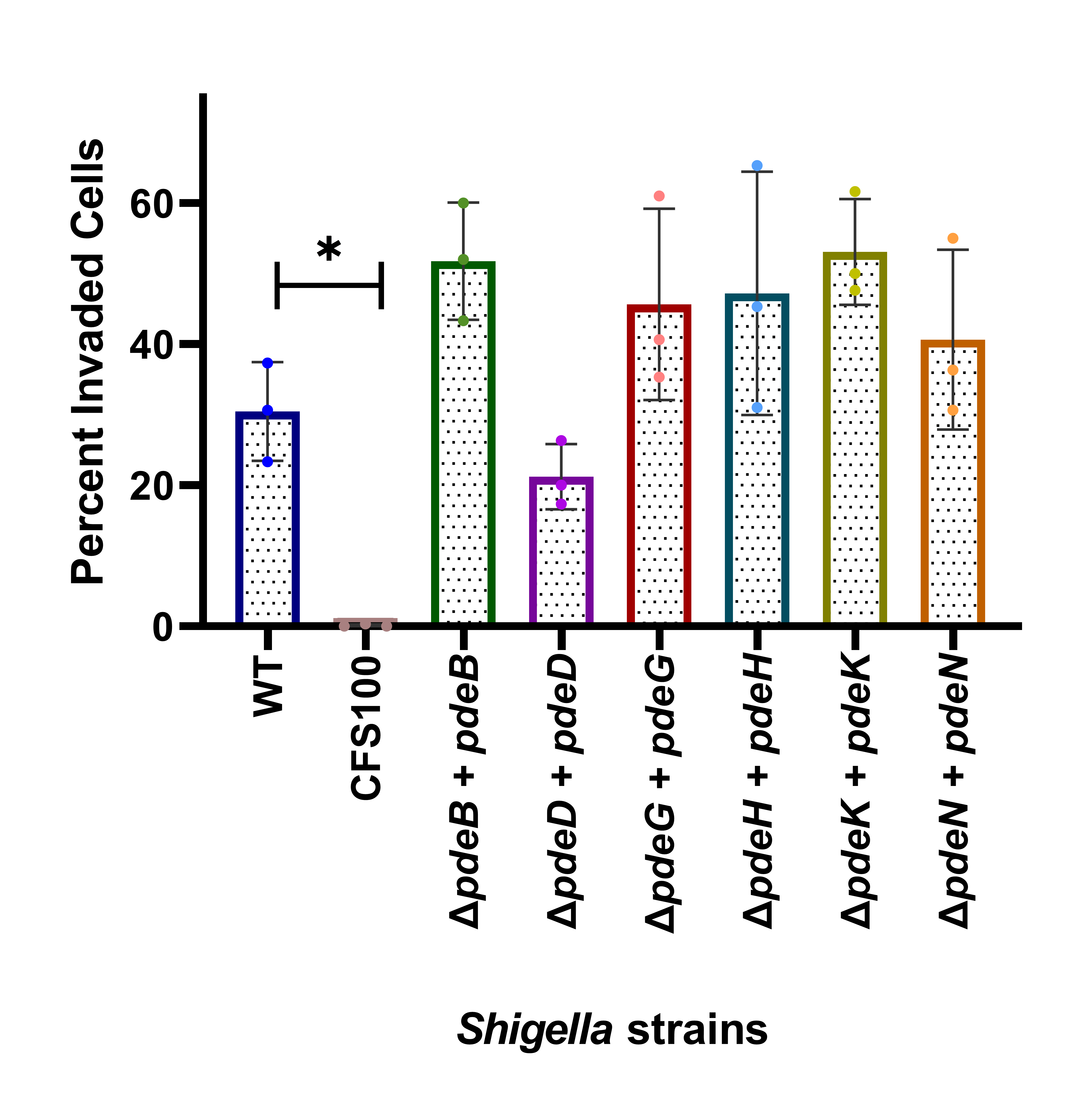

### Fig S4

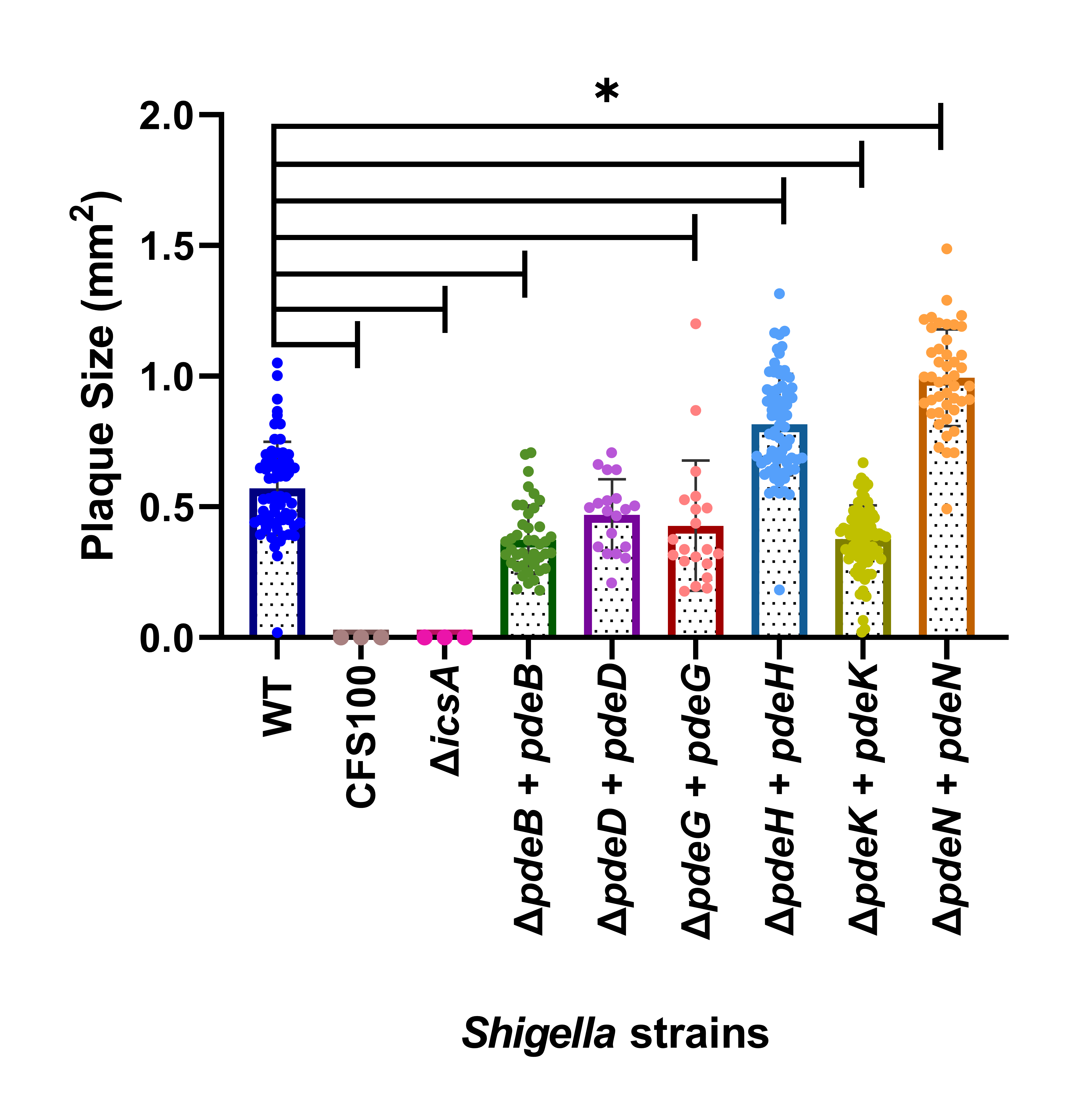

### Fig S5

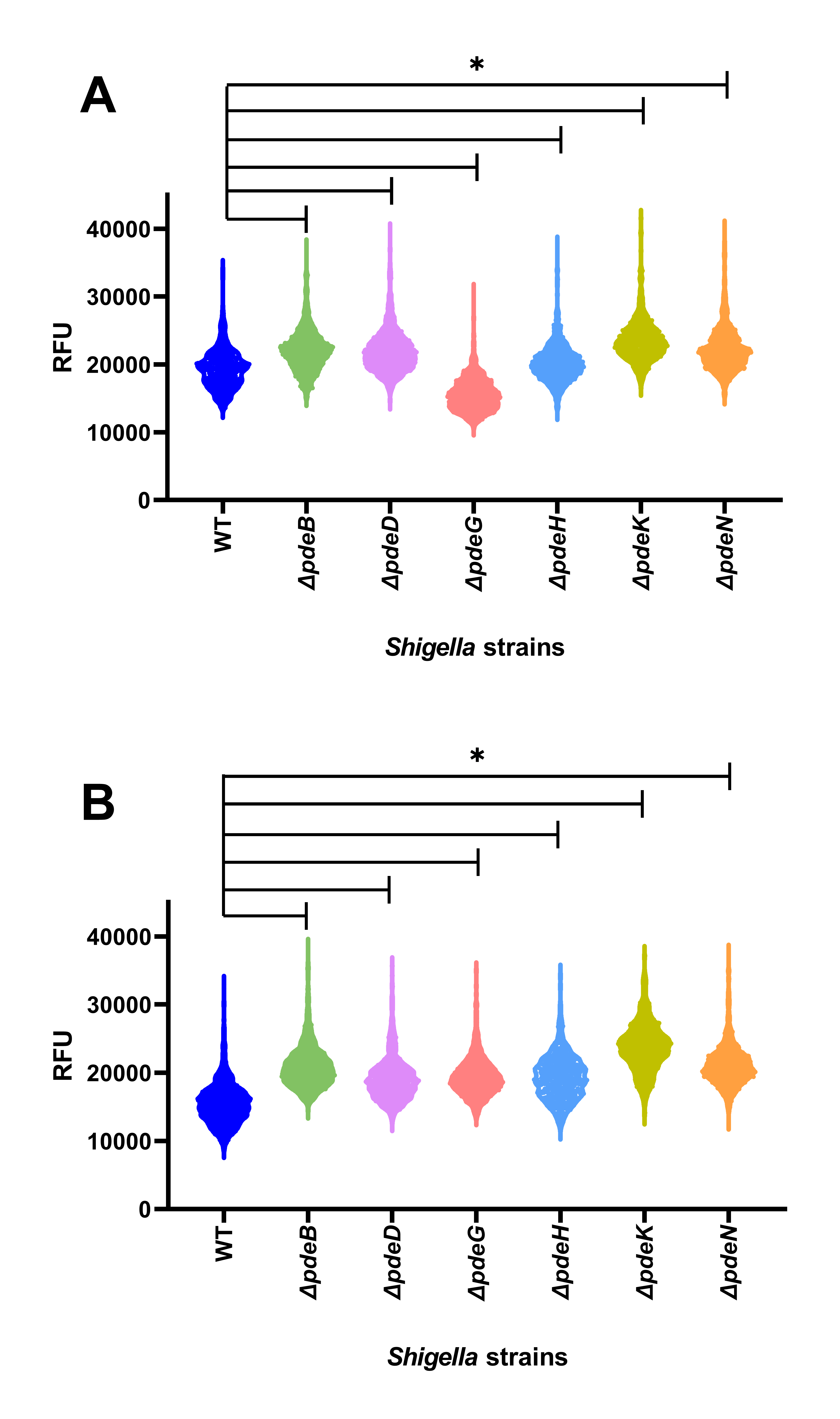

### Fig S6

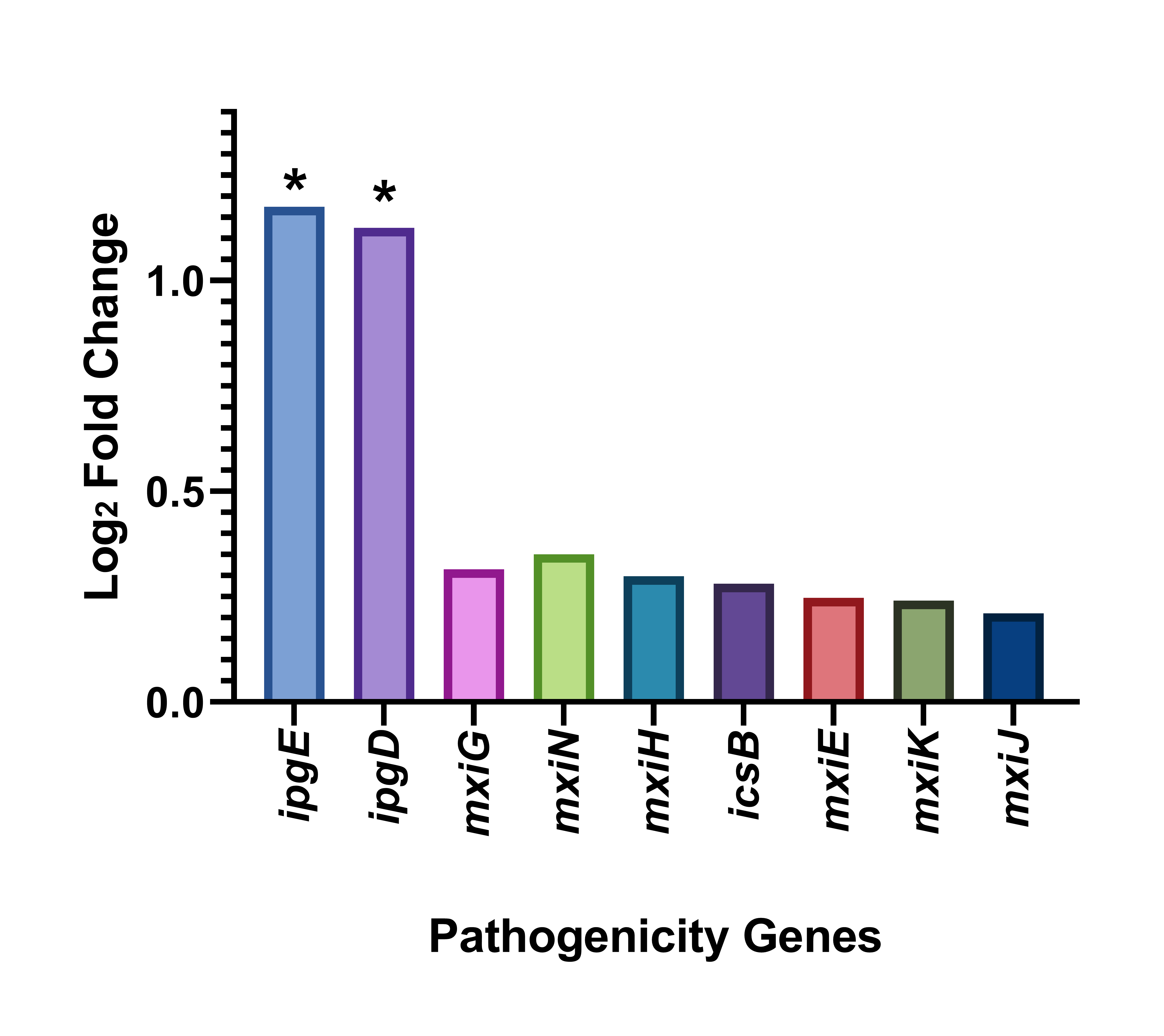

### Fig S7

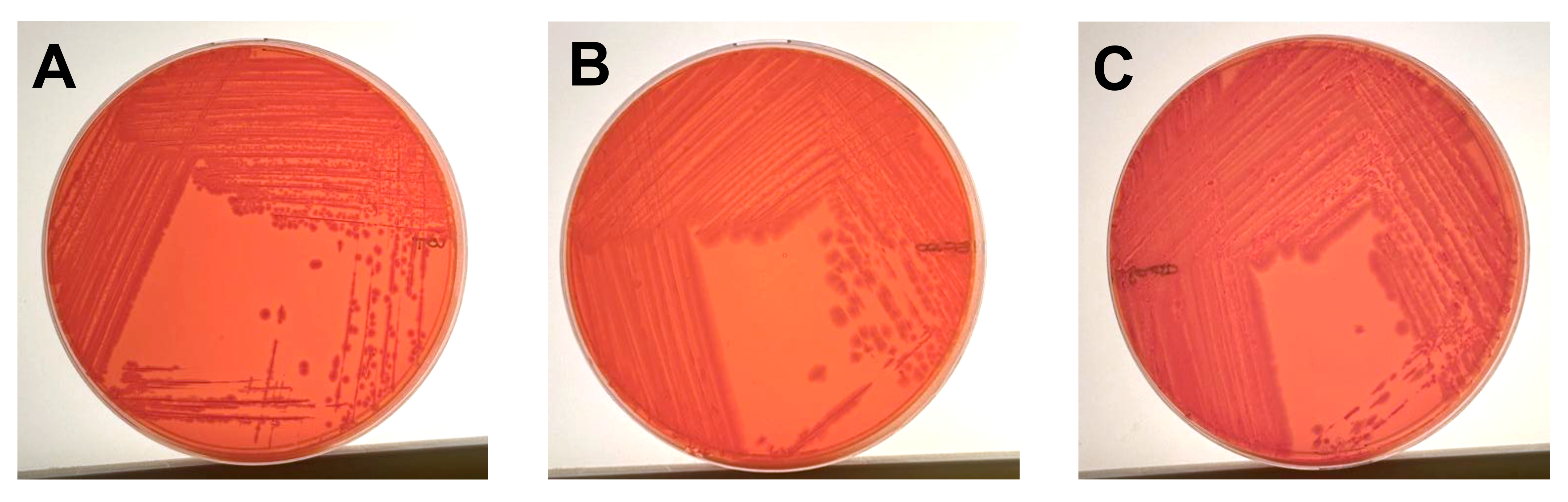

### Fig S8

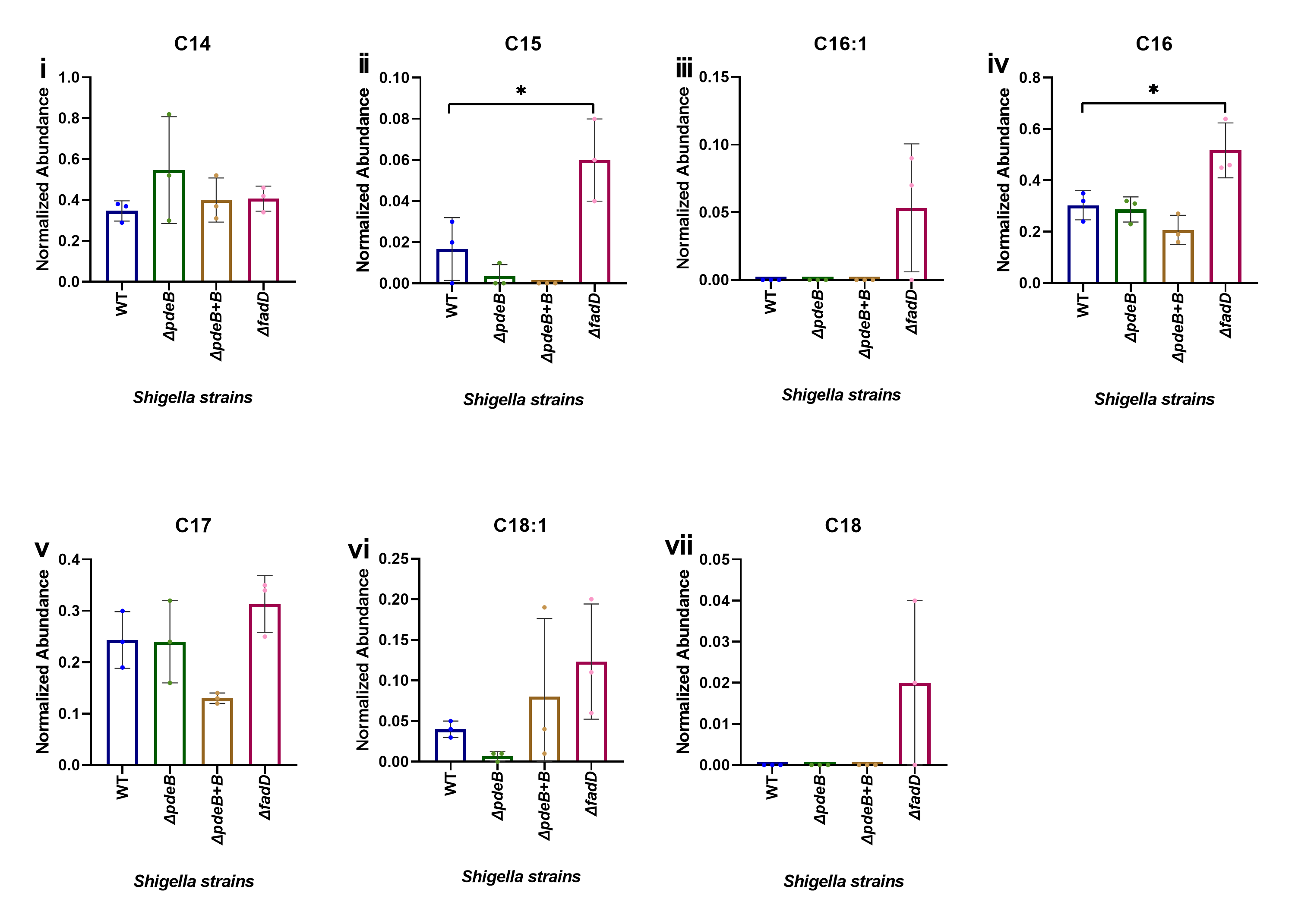
